# DNA methylation entropy is associated with DNA sequence features and developmental epigenetic divergence

**DOI:** 10.1101/2022.12.21.521278

**Authors:** Yuqi Fang, Zhicheng Ji, Weiqiang Zhou, Jordi Abante, Michael A. Koldobskiy, Hongkai Ji, Andrew P. Feinberg

## Abstract

Epigenetic information defines tissue identity and is largely inherited in development through DNA methylation. While studied mostly for mean differences, methylation also encodes stochastic change, defined as entropy in information theory. Analyzing allelespecific methylation in 48 human tissue sample datasets, we find that methylation entropy is associated with specific DNA binding motifs, regulatory DNA, and CpG density. Then applying information theory to 42 mouse embryo methylation datasets, we find that time- and tissue-specific patterns of development are more strongly correlated with methylation entropy than with mean, and methylation entropy is associated with sequence and chromatin features conserved with human. Moreover, methylation entropy is directly related to gene expression variability in development, suggesting a role for epigenetic entropy in developmental plasticity.

## Introduction

DNA methylation, a covalent modification of the nucleotide cytosine, heritable during cell division at CpG dinucleotides, is a key component of the epigenetic information, i.e. independent of the DNA sequence itself, defining cell type identity and developmental state. Differences in DNA methylation levels between individuals can be driven by nearby DNA sequence differences termed methylation quantitative trait loci (mQTLs) (1), and DNA methylation level is generally inversely related to mean gene expression levels, particularly at gene promoters (2). In addition to DNA methylation levels, DNA methylation stochasticity, more formally defined as entropy in information theory, is related to processes involving plasticity, such as the epithelial-mesenchymal transition in cancer (3) and differentiation potency, and could also help to regulate developmental plasticity (4). However, it remains poorly understood how epigenetic entropy is influenced by DNA sequence, how entropy might function, its relationship to developmental state or transcription factor binding sites, or what might be its effect on gene expression.

Information theory is the quantitative field that measures and analyzes information and entropy in its transmission, and it is thus a natural fit to address stochasticity in methylation. Figure 1A illustrates this idea applied to multiple consecutive CpG sites on the same DNA molecule. The two examples have similar mean methylation levels but markedly different entropy. The example on the right shows substantial variation from molecule to molecule in terms of the combinatorial configurations of the binary methylation states of multiple CpG sites in the same molecule, and hence high entropy; whereas the example on the left has little such variation and low entropy. Our overall goal was to relate differences in entropy (compared to mean) to the methylation potential energy landscape, genetic sequence, regulatory DNA, transcription factor binding, embryonic development, and gene expression (Figure 1A).

**Figure 1.**
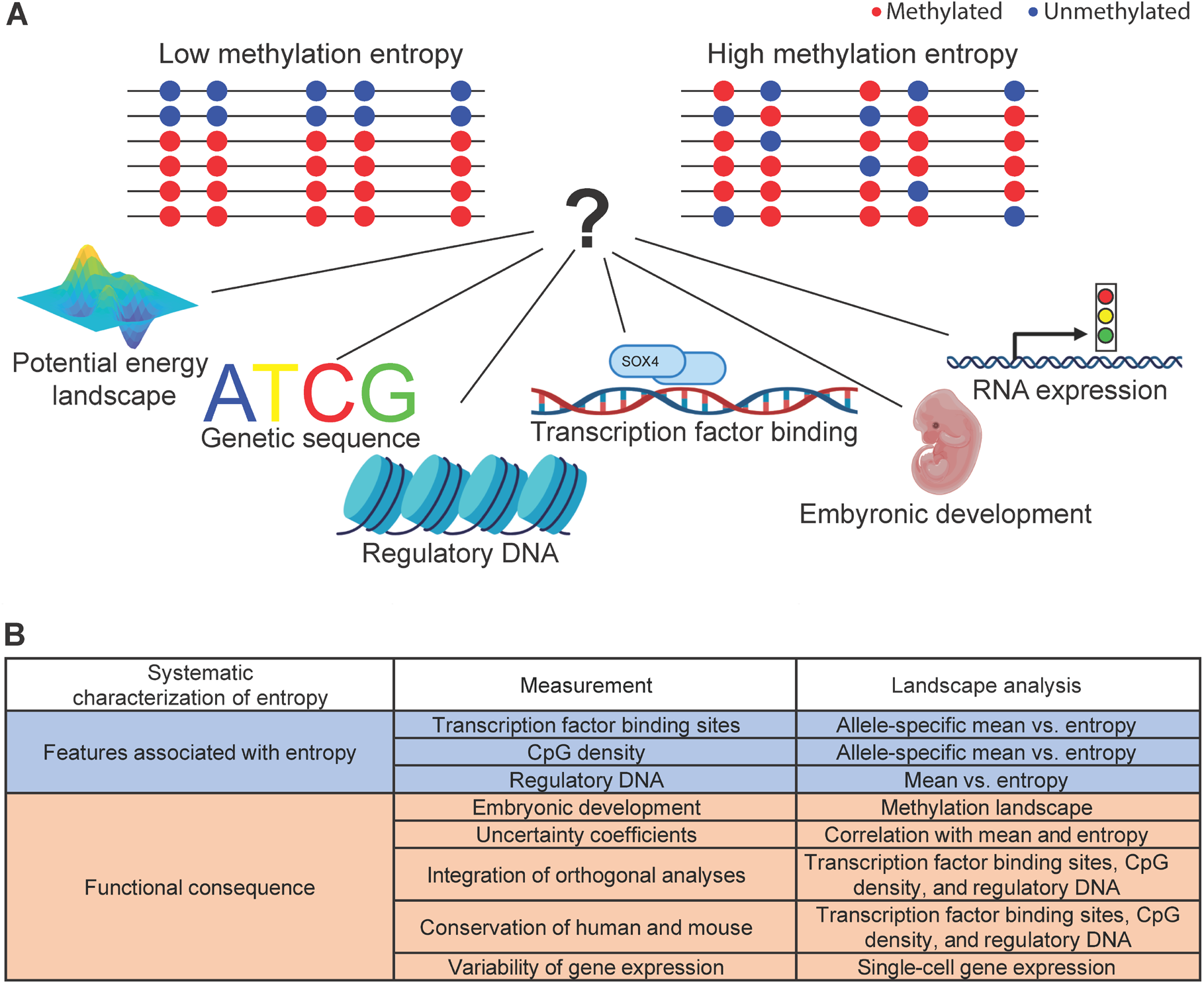
Study overview. **A.** Conceptual illustration of applying information theory to a set of DNA reads from WGBS. Each row is a single DNA molecule representing one read, and the dots on the line represents the CpG location. The blue dots correspond to unmethylated CpG and the red dots correspond to methylated CpG. The mean methylation level (MML) on the left panel is the same as the MML on the right panel. However, when considering the combinatorial configuration of methylation states of multiple CpGs in the same molecule, the left panel shows small variation among reads (rows), whereas the right panel shows large variation. As a result, the right panel has a much larger normalized entropy (NME) than the left panel (see Methods and Materials). **B.** A summary of the analyses performed in this study, which examines methylation entropy and its relationship to functional genomic features including transcription factor binding sites, CpG density and regulatory DNA; and relationship to functional consequences including the embryonic development, contribution of methylation entropy and mean methylation to methylation landscape, orthogonal relationships to transcription factor binding sites, CpG density and regulatory DNA, conservation of those features between human and mouse, and the association between gene expression variability and methylation entropy.

Entropy analysis has been applied to whole-genome bisulfite sequencing methylation data (5–7), but entropy analysis has not incorporated the role of the underlying DNA sequence itself in its control or been applied to embryonic development. We first applied an information theory method to measure DNA methylation mean and entropy that can distinguish the two alleles of a gene, providing the perfect control since the two alleles are present in exactly the same cells (8). We analyzed data from 49 polymorphic *human* samples obtained from the Roadmap Epigenomics Project, which contain not only DNA methylation sequencing data, but also complete DNA sequence (SNPs) (9). We asked three questions comparing entropy and mean methylation (outlined in blue in Figure 1B): (1) How are they related to transcription factor binding site? (2) How are they related to CpG density? and (3) How are they related to regulatory DNA?

In order to better understand the functional role of methylation entropy, we then performed an information-theoretic analysis of mouse embryonic development, examining comprehensive whole-genome DNA methylation data from the ENCODE3 project (10) on 7 individual tissue types during mouse development at 6 developmental time points, addressing five further questions (outlined in orange in Figure 1B): (1) Does this information theory-based approach identify tissue- and time-dependent changes in the methylation landscape? (2) What is the relative contribution of mean methylation and methylation entropy to these developmental landscape changes? (3) Can we integrate these results with orthogonal measures of transcription factor binding sites, CpG density, and regulatory DNA in order to understand the role of DNA methylation entropy? (4) Are these features conserved between mouse and human? and (5) What is the functional relationship between methylation entropy and gene expression variability, using singlecell RNA-seq data from both mouse and human? These analyses taken as whole revealed striking associations of methylation entropy with genomic and developmental processes, the sequences that may drive them, and their relationship to developmental gene expression.

## Materials and Methods

### SNP data processing

The revealing Single Nucleotide Polymorphisms (SNPs) information for subjects H9, HUES64, STL001, STL002, STL003, skin03, HuFGM02, 149,150, and 112 were extracted from the Roadmap Epigenomics database (11). Whole-genome sequencing data (WGS) for the H1 cell line was downloaded from PRJNA285681 (SRR2048232). The WGS reads were trimmed using Trim Galore (v0.5.0) (12). The trimmed reads were aligned to the hg19 reference genome using the Arioc WGS alignment package(v1.40) (13). Duplicated reads from polymerase chain reaction (PCR) products were removed using Picard tools (MarkDuplicates v2.18.13)(14). SNPs were called using GATK HaplotypeCaller (v4.0.0) (14) and dbSNP build 15 (15).

### Allele-specific methylation detection

The Whole-genome bisulfite sequencing (WGBS) data was aligned, and reads were assigned to each allele in the same way as the correlated potential energy landscape (CPEL) pipeline (8). The first and last 5 bp of each read were not used in the analysis. To perform haplotype-dependent allele-specific methylation analysis, we used the Julia package CpelAsm.j1, a recently developed method (8). For a given haplotype, CpelAsm estimates an allele-specific epigenetic landscape of the random methylation state ***X*** ∈ *χ* ≡ {0,1}^*N*^, where *N* is the number of CpG sites in the region, by performing maximum likelihood estimation on a set of *M* independent WGBS reads obtained for each allele as described in the CpelAsm (8), resulting in a probability mass function of the methylation state *p(**x**), x* ∈ *χ*, for each allele (allele subscript not shown for notational simplicity hence forth). Once the parameters of the allele-specific models are estimated, CPEL computes the allele-specific mean methylation level (MML) *μ* as well as the normalized methylation entropy (NME) *h*, a measure of epigenetic stochasticity, for each allele. In particular, the MML of a given allele is given by

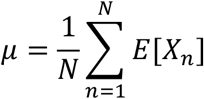

where *X_n_* is the random methylation state of the *n*-th CpG site, and *E*[*X_n_*] denotes the expected methylation level of the *n*-th CpG site. The corresponding NME is given by

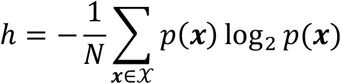

Both quantities are normalized to produce values in the range [0,1]. Then, CPEL computes the absolute MML difference and NME difference between two alleles. The first statistic measures the absolute difference in MML (dMML) between haplotype alleles, and it is simply given by |*μ*_1_ − *μ*_2_|. The second statistic measures the absolute difference in NME (dNME) between haplotype alleles, and it is given by |*h*_1_ − *h*_2_|. Thus, a large NME difference suggests that the methylation state in one allele is highly stochastic while that of the other allele behaves almost deterministically. Bootstrapping was used to perform hypothesis testing and compute a p-value for each statistic. Subsequently, CpelAsm used the Benjamini-Hochberg procedure to compute adjusted p-values to control the false discovery rate (FDR) in the statistical output (16). The FDR ≤ 0.1 was used to determine the significance of the MML difference and NME difference.

### Regions with large MML difference enrichment analysis

The list of imprinted genes was downloaded from the Geneimprint website (17). The contingency table was constructed, by counting the occurrence that the regions showing large MML differences in the promoter of the imprinted gene versus not in the promoter of the imprinted gene. The odds ratio was calculated using the two-sided Fisher’s exact test from the contingency table. The same enrichment analysis was done for the mono-allelic expressed gene (MAE) (18).

### Human motif binding analysis

We used the motifBreakR package in the default setting to first map the motif obtained from the JASPAR database (19) at each SNP to predict the transcription factor binding sites (TFBSs). For each motif site, we then evaluated the binding probability of the corresponding motif in each allele by considering the polymorphism. For each motif site, we counted the occurrence of the region with higher binding probability in the allele with higher NME or MML than the other allele. This observation should be 50% by random chance. On the contrary, if the occurrence of such regions is significantly higher than 50%, using the binomial test, it suggests that a higher transcription factor (TF) binding probability is associated with higher NME or MML. Similarly, we performed the analysis to identify motifs that are associated with lower NME or MML.

### Analyzing the association between NME and SNP

We examined the trinucleotide context near the SNP, i.e. 1bp before and after the SNP. For each type of SNP, there are 16 possible trinucleotide changes. We merged trinucleotide changes that are reverse complements, e.g., GCG → GTG and CGC → CAC. The trinucleotide was connected by → in the direction of SNP. The direction of SNP was determined by decreasing CG if there are CG changes in the trinucleotide, e.g., GCG → GTG. Allele1 is the allele on the left, e.g., GCG, and the allele2 is the allele on the right, e.g., GTG. The NME difference was calculated for each region using NME in the allele1 minus the NME in allele2. For each trinucleotide context, the contingency table was constructed by counting the occurrence that the NME in allele1 is smaller than the NME in allele2 in that trinucleotide context versus not in that trinucleotide context (Supplementary Figure S5). The log (Odds ratio) and p-value were calculated using the two-sided Fisher’s exact test. The Benjamini-Hochberg correction was used to calculate the FDR for significance (16).

Some regions contain more than 1 SNPs that change the CpG number. To calculate the effect of the CpG number in the allele on NME, we calculated the CpG number changes and NME changes between two alleles. The analyzed regions with different CpG numbers between two alleles due to SNP and significant NME difference (FDR ≤ 0.1) were selected. Among those regions, the distribution of NME in the allele with more CpGs was plotted in Figure 3B in red and the distribution of NME in the allele with fewer CpG was plotted in Figure 3B in green. The one-sided t-test was used to test the significance NME difference between two alleles because we found the allele with more CpG tends to have smaller NME when analyzing SNPs. The NME in both alleles in all the regions with significant NME differences between the two alleles, regardless of CpG number, was plotted in Figure 3B in blue as a reference.

**Figure 2.**
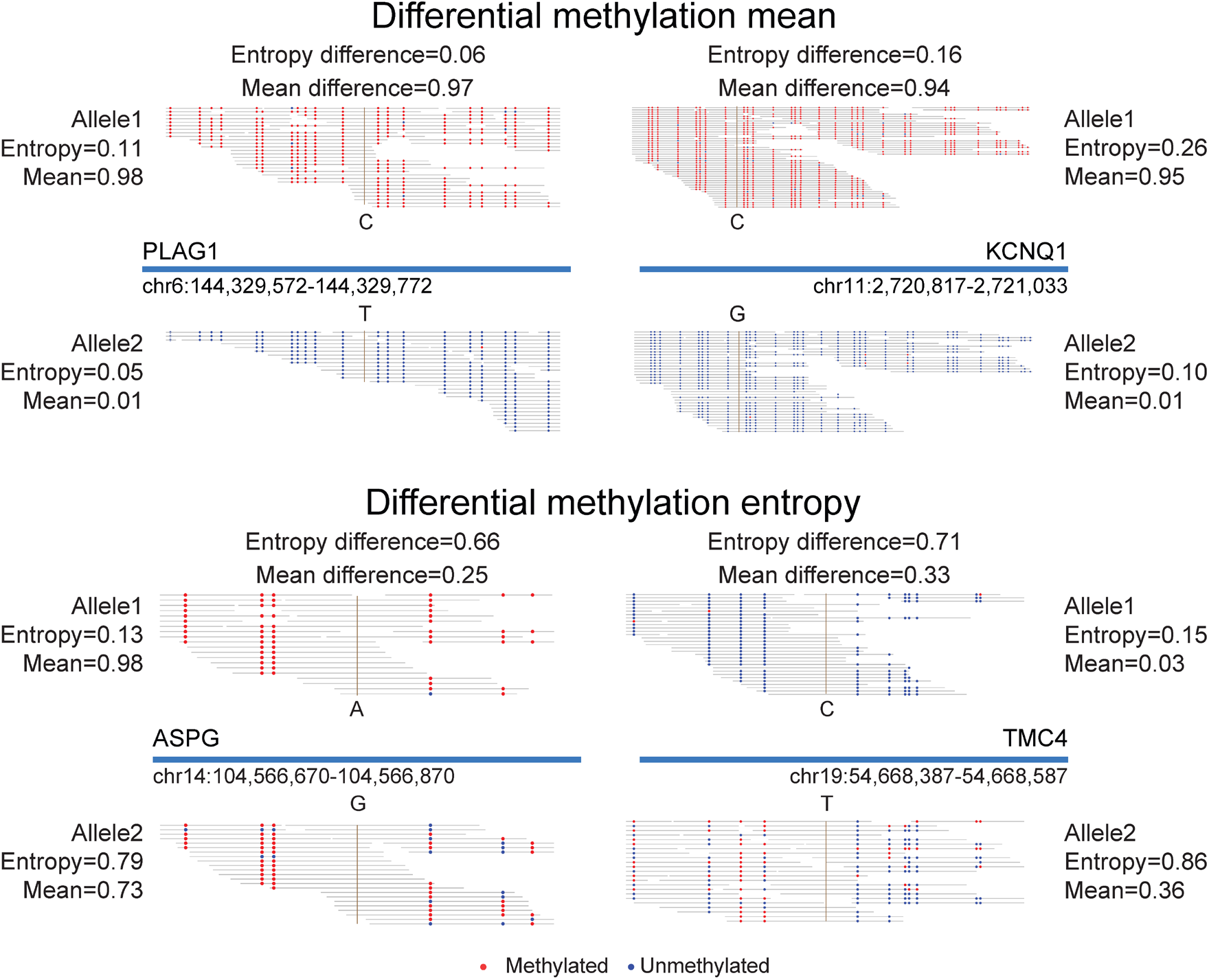
Examples of allele-specific methylation mean and entropy analysis. Four example regions with large methylation mean difference (top panel) or large methylation entropy difference (bottom panel) between two alleles. For each region, the continuous grey horizontal lines represent individual sequencing reads. The brown vertical lines are the location of the SNPs used to distinguish two alleles. The red dots represent methylated CpG sites, and the blue dots represent unmethylated CpG sites. The top panel shows two example regions located at two known imprinted genes, *PLAG1* and *KCNQ1*. Consistent with the known association between imprinting and promoter methylation, in one allele almost all CpG sites are methylated and in the other allele almost all CpG sites are unmethylated. The lower panel shows two genes, *ASPG* and *TMC4*, with high methylation entropy differences inside the gene. For *ASPG*, the allele with genotype A has low entropy (0.13) as indicated by most of the reads having the same methylation status. The allele with the genotype of G has high entropy (0.79) as indicated by the highly stochastic color pattern. For *TMC4*, the allele with the genotype of C has low entropy (0.15), while the allele with the genotype of T has high methylation entropy (0.86) as indicated by the highly variable methylation patterns across reads.

**Figure 3.**
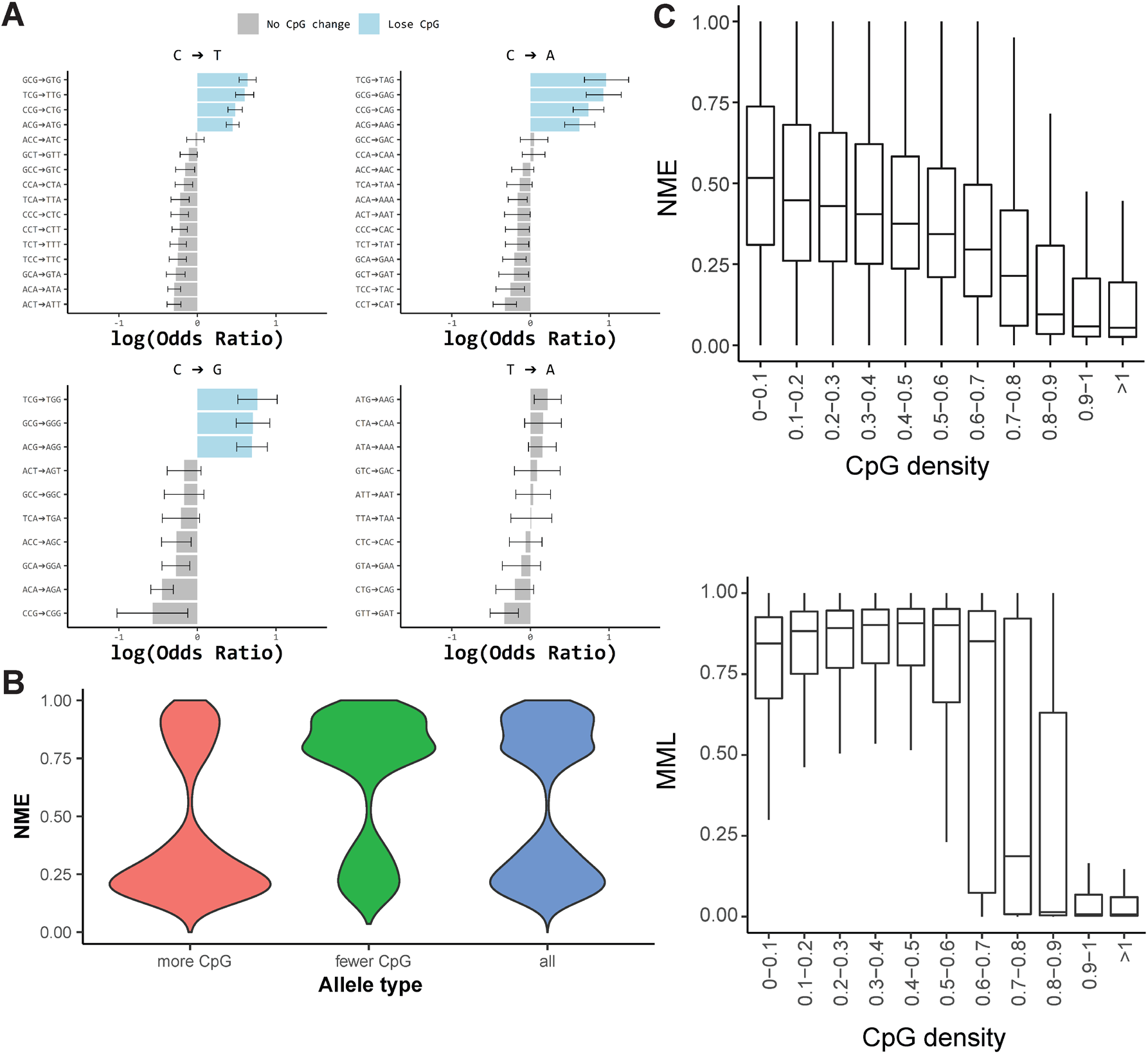
Allelic sequence relationship to NME. **A.** The analysis for each SNP genotype in all 52 possible trinucleotide contexts. Log(Odds ratio) (x-axis) greater than 0 means the right trinucleotide is more likely to have higher entropy than the left trinucleotide than random expectation. If the SNP changes the number of CpG sites, the allele with fewer CpG sites is shown on the right and the bar is colored in blue. For each SNP genotype, the trinucleotide contexts associated with the largest allelic entropy differences (i.e. highest odds ratio) were often the ones that change the CpG number (colored in blue). Error bar is used to show the upper and lower 95% of CI of the log(OR). **B.** Distribution of NME comparing the alleles with higher or lower numbers of CpG. For the regions that have CpG number differences, the allele is labeled as “more CpG” (red) for the allele that contains more CG than the allele labeled “fewer CpG” (green). The allele with more CG has significantly lower NME than the allele with fewer CpG (paired one-sided t-test, *P* < 2.2 × 10^-16^). The distribution of NME for all alleles is displayed as a reference (blue). **C.** NME and MML, regardless of SNPs, of genomic regions surrounding transcription start sites (20kb around each TSS) stratified based on CpG density. The CpG density was defined as the ratio of the observed CpG number to the expected CpG number in each region (see Methods and Materials). The centerline represents the median and the upper and lower lines are the first and third quantile. As CpG density increases, the NME decreases. MML showed an abrupt decrease at high CpG density while NME had a gradual decrease as CpG increased.

The number of CpG was counted in each allele with an extended region (500 bp). The expected number of CpG was calculated using equations from (20)

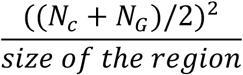

The density was calculated for each allele by the number of CpG/expected CpG. The CpG density difference was calculated by the difference of CpG density between two alleles. The regions showing significant NME difference (FDR ≤ 0.1) were used to calculate the correlation between CpG density difference and NME difference.

### Human NME and MML calculation without separating alleles

Allele-specific analysis can only be performed for regions with heterozygote SNPs. To analyze the whole-genome regardless of SNPs, the WGBS data for human samples were aligned without assigning the allele. The first and last 5 bp of each read were not used. The target region was divided into 250 bp segments. For each segment, regardless of the allele of origin, the MML and the NME were calculated as described in the **Allele-specific methylation detection** section. The CpG density-dependent nature of differential NME (dNME) suggested that simply lumping data from both alleles might give a false impression of information-theoretic entropy. If one simply pooled the reads from the two alleles with a 50-50 mixture of a highly methylated and lowly methylated allele of a gene, both with near-zero entropy, combined reads would sometimes falsely appear to be relatively high entropy. That is exactly what would happen when examining imprinted regions, where one allele is in fact methylated depending on the parent of origin. Indeed, if one computes NME without regard to an SNP, there is a false skewing to high NME in regions with significant MML difference such as genes with imprinting (Supplementary Figures S1A, B); while that is not the case for regions without significant MML difference (Supplementary Figure S1C). Since regions with allelic mean methylation imbalances were rare across the genome (0.422% of all regions analyzed), and since the main difference between the mean allelic methylation entropy and methylation entropy regardless of SNP were in such regions, we excluded regions showing mean methylation imbalances from this analysis. As a result, in this whole-genome analysis, NME is expected to be similar to the mean allelic NME of the two alleles at each locus.

### NME and CpG density analysis

NME and MML were calculated within 20 kb of transcription start site (TSS) as described in the **Human NME and MML calculation** section. We computed the CpG density for each region using (total number of CpG)/(expected number of CpG). The expected number of CpG was calculated using the same way we did in **Analyzing the association between NME and SNP**. The correlation between CpG density and NME or MML regardless of the allele was calculated using the two-sided Pearson correlation test.

### Human motif analysis

We downloaded the non-redundant CORE motifs from JASPAR (19). The 630 human motifs representing 586 TFs or TF complexes were then mapped to the human genome using CisGenome (21). For each motif, the mapped motif sites were grouped into two classes (regulatory and non-regulatory) based on 167 ENCODE (22) DNase-seq samples. Regulatory DNA is defined as the union set of chromatin accessible sites across the 167 DNase-seq samples which were downloaded from https://github.com/WeiqiangZhou/BIRD-data (23). To get a set of non-regulatory DNA, we used CisGenome (21) to obtain genomic regions that are not located in the regulatory DNA but have a similar distribution of distance to TSS with the regulatory DNA. For each motif, motif sites that overlap with regulatory DNA were labeled regulatory motif sites, and motif sites that overlap with non-regulatory DNA were labeled as non-regulatory motif sites. For this, we used methylation entropy to characterize the NME of each motif site in regulatory DNA and non-regulatory DNA. The NME regardless of alleles was calculated using the method described in the Human **NME and MML calculation** section. For each TF, the median value of NME across all motif sites in each of the two motif site classes (i.e. regulatory and non-regulatory) was computed. Based on the median NME for each TF across different samples, we then used a two-sided Wilcoxon signed-rank test to test whether NME is different between the two classes of sites. To adjust for multiple testing, we calculated the false discovery rate (FDR) using Benjamini–Hochberg procedure (16). Supplementary Figure S2A shows the standardized NME values where the median NME for each TF was standardized to have zero mean and unit standard deviation across all samples and the two classes of sites. Similarly, we compared the MML between motif sites in regulatory DNA and non-regulatory DNA for each TF. Supplementary Figure S2D shows the standardized MML values based on the median MML for each TF. We also applied a two-sided Wilcoxon signed-rank test to test whether MML is different between the two classes of sites.

### Mouse embryonic development data analysis

The aligned WGBS bam files were manually downloaded from the ENCODE3 database. The bam files were sorted using samtools v1.9 and deduplicated using Picard tools MarkDuplicates (v2.23.3-4) function. Uncertainty coefficient (UC) is the Jensen-Shannon distance (JSD) normalized by entropy between the DNA methylation landscapes of two time-points which captures both mean methylation changes and methylation entropy differences. We quantified the difference in methylation landscapes between the two samples *s*_1_ and *s*_2_ by computing the uncertainty coefficient (UC)

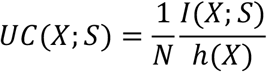

where *I*(*X;S*) is the mutual information between the methylation state X and the sample S, and it is given by

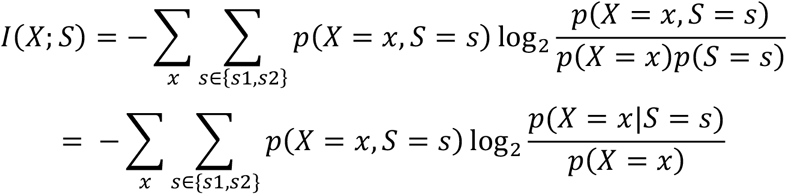

and h(X) is the NME under a model where the sample S has been integrated out. It has been shown that *I*(*X; S*) = *D*^2^(*p*_1_, *p*_2_) where *D*(*p*_1_, *p*_2_) is the Jensen Shannon Divergence (JSD) that measures the similarity between two probability distributions *p*_1_ = *p*(*X* = *x*|*S* = *s*_1_) and *p*_2_ = *p*(*X* = *x*|*S* = *s*_2_ (8). Large JSD means a large difference in methylation probability distribution between two samples. Given the linear dependency of *UC*(*X;S)* on the mutual information *I(X;S)*, large values of *UC(X;S)* imply a large difference between samples. The NME, MML, differential NME, differential MML, and UC were then calculated using the CPEL pipeline for these genomic regions (8). Regions that have lower than 10x coverage were excluded.

For each tissue type, regions with UC > 0.1 were first analyzed for clustering analysis. Because we found more than 28% of them only have UC > 0.1 in one tissue, we selected the region with UC > 0.1 only within that tissue (not in other tissues) for at least one pair of time points. UC for all consecutive pairs of time points (e.g., E10.5 and E11.5, E11.5 and E12.5) were standardized to have mean 0 and variance 1 for each region across consecutive pairs of time points. The standardized values were then used to computationally group regions into 10 UC clusters with k-means clustering. The k-means clustering was performed 10 times with different random seeds. The regions that were always clustered in the same clusters were defined as core-cluster regions. The rest of the regions were assigned to each core cluster based on their correlation to the mean UC of the core cluster. The 10 UC clusters were ordered according to which consecutive time points have the largest UC.

We selected UC > 0.1 as the cutoff because the top 5% of all UC values is around 0.1. We further tested the other UC cutoffs, 0.025 (~top 30%), 0.05 (~top 15%), 0.15 (~top 3%), 0.2 (~top 1%), using the same method.

### Mouse gene ontology (GO) analysis

Mouse GO analysis was done for each cluster in each tissue. For regions overlapping with enhancers, the target genes were annotated according to the published dataset (24). The genes for which we got complete methylation data in all samples at enhancer regions were used as the background gene list for GO analysis. For each cluster, the regions overlapping with enhancers were annotated to the corresponding gene and were used as target genes for GO analysis. The R package topGO was used to perform GO analysis using the classicFisher method (25). The GO terms with more than 10 annotated genes were retained. The GO terms that have 0 significant genes were excluded. The p-values were calculated conditional on the knowledge that the GO terms with 0 significant genes were excluded. The Benjamini–Hochberg procedure was used to correct p-values for multiple hypothesis testing (16). The GO terms with fold change (FC) greater than 1.5 were reported. The final GO terms for each cluster were ranked by FDR and FC was used to break ties. The top 5 GO terms were selected from each cluster to generate the heatmap. The GO terms for all tissues and clusters were plotted together. The GO terms that do not have FDR ≤ 0.2 in all tissue-cluster were excluded. The rows in the heatmap were ordered based on which columns show the highest FC.

For regions overlapping promoters, defined as within 2kb of TSS, the genes were annotated to that TSS and the GO analysis was performed using the same procedure. The genes for which we have complete data in all samples at promoter regions were used as the background list.

We further tested the other UC cutoffs, 0.025 (~top 30%), 0.05 (~top 15%), 0.15 (~top 3%), 0.2 (~top 1%), using the same method.

### Categorize regions based on UC-dNME and UC-dMML correlation

For each tissue, we calculated the correlation between NME change and UC (dNME-UC correlation) and the correlation between MML change and UC (dMML-UC correlation). We randomly permuted the labels of the NME/MML differences 20 times, recalculated the correlations between permuted NME/MML and UC as the null distribution, and obtained empirical p-values as the tail areas under the null distribution. FDRs were then calculated based on p-values using the Benjamini–Hochberg procedure (16). Some regions have a high dMML-UC correlation and dNME-UC correlation (Supplementary Figure S3A). To distinguish those regions from the regions that only have high dNME-UC correlation or dMML-UC correlation, we identified different clusters of regions based on the density of dMML-UC correlation and dNME-UC correlation. To that end, we calculated the difference between dNME-UC correlation and dMML-UC correlation as well as the mean of two correlations for each region (Supplementary Figure S3B). We binned the mean correlation with a 0.05 interval. For each bin, we search for the first local minimum for the density of dNME-UC and dMML-UC difference below 0 and above 0 (negative local minimum and positive local minimum). The regions with values below the negative local minimum suggested the dMML-UC correlation is larger than the dNME-UC correlation (Supplementary Figures S3C, D). If those regions also have significant dMML-UC correlation (FDR ≤ 0.2) or have dMML-UC correlation greater than 0 while dNME-UC correlation is smaller than 0, they were categorized as regions whose UC is predominantly correlated with MML change (predominantly MML-correlated regions), meaning that their UC changes across time can be mainly explained by the MML changes (Supplementary Figure S3E, F). On the other hand, the regions with values above positive local minimum suggested the dNME-UC correlation is larger than the dMML-UC correlation. If their dNME-UC correlation is significant (FDR ≤ 0.2) or their dNME-UC correlation is greater than 0 and dMML-UC correlation is smaller than 0, they were categorized as regions whose UC is predominantly correlated with NME change (predominantly NME-correlated regions), meaning that their UC changes across time can be mainly explained by the NME changes (Supplementary Figure S3E, F). The regions that have values between two local minimums and have significant dNME-UC correlation and dMML-UC correlation (FDR ≤ 0.2) were categorized as regions whose UC is correlated with both NME and MML, meaning that both MML and NME changes contribute to the temporal changes of UC (Both). The regions that have insignificant dMML-UC correlation and dNME-UC correlation were categorized as regions whose UC is independent of both NME and MML (Neither) (Supplementary Figure S3E, F). The one-sided t-test was used to determine if there are more regions whose UC is predominantly correlated with NME change than regions whose UC is predominantly correlated with MML change. We chose a one-sided t-test because visually there are more regions whose UC is predominantly correlated with NME than regions whose UC is predominantly correlated with MML. We did the same analysis using the other UC cutoffs, 0.025, 0.05, 0.15, 0.2.

### Mouse motif analysis

We downloaded 736 human and mouse motifs from JASPAR and mapped the motifs to the mouse genome using CisGenome (21). Enrichment of each motif was calculated as the ratio between the odds of motif sites in the target regions (i.e. [number of target regions that contain motif sites]/[number of target regions that do not contain motif sites]) and the odds of motif sites in control regions (i.e. [number of control regions that contain motif sites]/[number of control regions that do not contain motif sites]). The control regions were obtained using CisGenome (21) which has a similar distribution of distance to TSS with the target regions. One-sided Fisher’s exact test was applied to test whether the motif is significantly enriched in the target regions. Multiple testing was adjusted by converting P-values to FDRs using Benjamini–Hochberg procedure (16). To compare the enrichment of each motif between regions where the methylation landscape change (i.e. UC) is predominantly correlated with NME with regions where UC is predominantly correlated with MML, we fitted a linear regression model to the normalized log odds ratio of motif enrichment from the two types of regions. The normalized log odds ratio was calculated using the log odds ratio divided by its standard error. We identified TF motifs that were outside the 75% prediction interval and were significantly enriched in either region type (FDR ≤ 0.1). TF motifs that were more enriched in regions where UC is predominantly correlated with NME were marked with red color and defined as “entropy-associated motifs”, and TF motifs that are more enriched in regions where UC is predominantly correlated with MML were marked with blue color and defined as “mean-associated motifs”.

We also compared the entropy-associated TF motifs combined from all tissue types with TF motifs preferring high NME in the allele-specific analysis in human. By performing a one-sided Fisher’s exact test, we found that compared to mean-associated TF motifs, entropy-associated TF motifs in mouse have significantly higher overlap with the human high NME-allele preferring motifs. The one-sided Fisher’s exact test was used because the assumption is the entropy-associated TF is more enriched in mouse than mean-associated TF. Similar to the analysis in humans, we studied the relationship between NME and regulatory DNA in the mouse. We compared the NME between motif sites that are located in regulatory DNA as defined by a union set of mouse DNase I hypersensitive sites and those located in non-regulatory DNA (obtained similar to the human analysis) in each mouse sample for 736 transcription factor binding motifs.

### Single-cell RNA-seq data processing

Human single-cell RNA-seq raw count data were downloaded from Human Cell Landscape (26) (Microwell-seq platform). Cells with at least 500 expressed genes with non-zero read counts were retained. The read counts were normalized by library size and gene expression values were imputed using SAVER (27) to address the high sparsity.

Mouse single-cell RNA-seq processed log2-transformed FPKM data were downloaded from ENCODE (28) (Fluidigm C1 SMART-seq platform). The log2-transformed expression matrix was transformed back to the original scale and imputed using SAVER (27).

For both human and mouse data, SAVER imputed values were then log2-transformed. Genes with non-zero expression in at least 10% of all cells were retained, and all ribosomal genes were removed. The processed data were used for the subsequent gene expression variability analysis.

### Calculation of gene expression mean-adjusted variability (MAV)

Let *y_ij_* be the library-size normalized, imputed, and log2-transformed expression level for gene *i*(*i* = 1,…, *I*) and cell *j*(*j* = 1,…, *J*). Let *m_i_* and *s_i_* be the mean and standard deviation of the expression level for gene *i* across all cells respectively. A B-spline regression model was fitted across all genes where *s_i_* is the response variable and *m_i_* is the independent variable, and let *ŝ_i_* be the fitted values of the standard deviations. The gene expression mean-adjusted variability (MAV) of gene *i*, *h_i_*, is defined as the residual of the regression model, or equivalently the difference between observed and fitted standard deviation: *h_i_* = *s_i_* – *ŝ_i_*.

## Results

### Methylation entropy can depend on DNA sequences

To identify DNA sequences specifically associated with methylation entropy, we applied our recently developed information-theoretic method for allele-specific methylation analysis(8) to 49 human samples from the Roadmap Epigenomics Project (9). This approach allows us to rigorously analyze genetic sequence-driven differences in methylation *in the exact same cellular and tissue context*. We used a methylation potential energy landscape model that considers all potential methylation states, cooperative interactions between adjacent sites, and adheres to the rigorous definition of Shannon entropy(8) (see Methods and Materials). In Figure 2, we illustrate four sets of sequencing reads, where the two alleles are distinguished by a single nucleotide polymorphism (SNP), from which we calculated mean methylation level (MML) and normalized methylation entropy (NME), i.e. normalized for the number of methylatable CpG sites. On the top are two genes, *PLAG1* and *KCNQ1* showing large mean differences, where one allele is much more methylated overall than the other. Explaining the large methylation difference between alleles, both *PLAG1* and *KCNQ1* are imprinted genes, i.e. with the parent of origin-specific expression(29,30), consistent with the known role of methylation in silencing imprinted loci on one allele(31,32). Indeed, we found that regions with large mean allelic methylation difference were enriched in imprinted genes (Supplementary Figure S4, Fisher’s exact test, OR= 60.5, 95% CI = [47.3 – 76.9], P < 2.2 × 10^-16^, Methods). Note that allelic differences of methylation in imprinted genes are *not related* to the underlying DNA sequence, the focus of the present study, because what is a maternal allele in one generation can be a paternal allele in the next. In contrast, at the bottom of Figure 2 are examples of large *sequence-driven* differential methylation entropy between individual alleles at *ASPG* and *TMC4*, with much smaller differences in mean methylation levels than the examples on top. Note that previously *sequence-independent* entropy was studied in a way that does not separate two alleles, and such analysis identified imprinted genes as in our examples in Figure 2 as if they had high methylation entropy even though they may not at allelic level (11). For example, for regions with the allelic difference in mean methylation like those in the top panel of Figure 2, if one mixes the two alleles, then the entropy may appear to be higher in the *allelic mixture*, e.g., 0.28 for PLAG1, than in the individual alleles, but at the sequence level, this increase in entropy is actually due to parent of origin-specific imprinting. For each allele, the entropy is low. In examples such as the bottom of Figure 2, the differences in entropy are specific to the DNA sequence (i.e., the entropy is high within an allele), which is our focus as we wish to understand the underlying sequence drivers of entropy. Among the 3,332,744 regions containing heterozygous SNPs we analyzed, 29,681 exhibited significant allele-specific differential NME (*FDR* ≤ 0.1), but only 6,807 regions exhibited significant differential MML (*FDR* ≤ 0.1).

### Entropy-associated DNA sequences are associated with predicted transcription factor binding sites

We next scanned the nucleotide sequence around each SNP for transcription factor binding sites (TFBS) based on motifs from the JASPAR database (19) and computed the binding probability in each allele as described (33). We identified 129 motifs with higher binding probability in the allele with a lower mean methylation level than the other allele, and only 7 motifs with higher binding probability in the allele with a higher mean methylation level than the other allele (Supplementary Dataset S1). Supporting the validity of our approach, CTCF was ranked 5th among the 129 motifs associated with low methylation levels (Supplementary Dataset S1, Method), consistent with the known observation that the allele with higher CTCF binding probability shows lower methylation (34). Similarly, 89%-93% of motif sites of NFI family members, including NFIC, NFIB, NFIX, showed lower mean methylation level in the allele with higher binding probability (Supplementary Dataset S1) than the other allele, consistent with the observation that the NFI family proteins are enriched at demethylated sites during neural development (35).

Using a similar approach, we then asked what motifs are associated with allelic differences in methylation entropy. We found 135 motifs with higher binding probability in the allele with significantly higher methylation entropy than the other allele (binomial test, *FDR* ≤ 0.1; Supplementary Dataset S2), compared to 14 motifs showing decreased binding probability in the allele with significantly higher methylation entropy at the same FDR level (binomial test, *FDR* ≤ 0.1; Supplementary Dataset S2). Among these 135 higher entropy-associated motifs, 55 were not associated with low MML (Table 1, Supplementary Dataset S3). Moreover, gene ontology (GO) enrichment analysis (36) of the transcription factors associated with these 55 motifs shows enrichment for positive regulation of cell development (enrichment = 5.43, *P* = 4.72 × 10^−5^) and positive regulation of cell differentiation (enrichment = 3.32, *P* = 7.71 × 10^−4^). Furthermore, several of the transcription factors associated with these motifs are pioneer transcription factors, i.e. that open up condensed chromatin, including *ASCL1, PBX1, MEIS1, ATF4, ESRRB*, and *KLF4* (37–42). In contrast, low MML-associated motifs not associated with high NME showed no GO enrichment of the associated transcription factors.

**Table 1.**
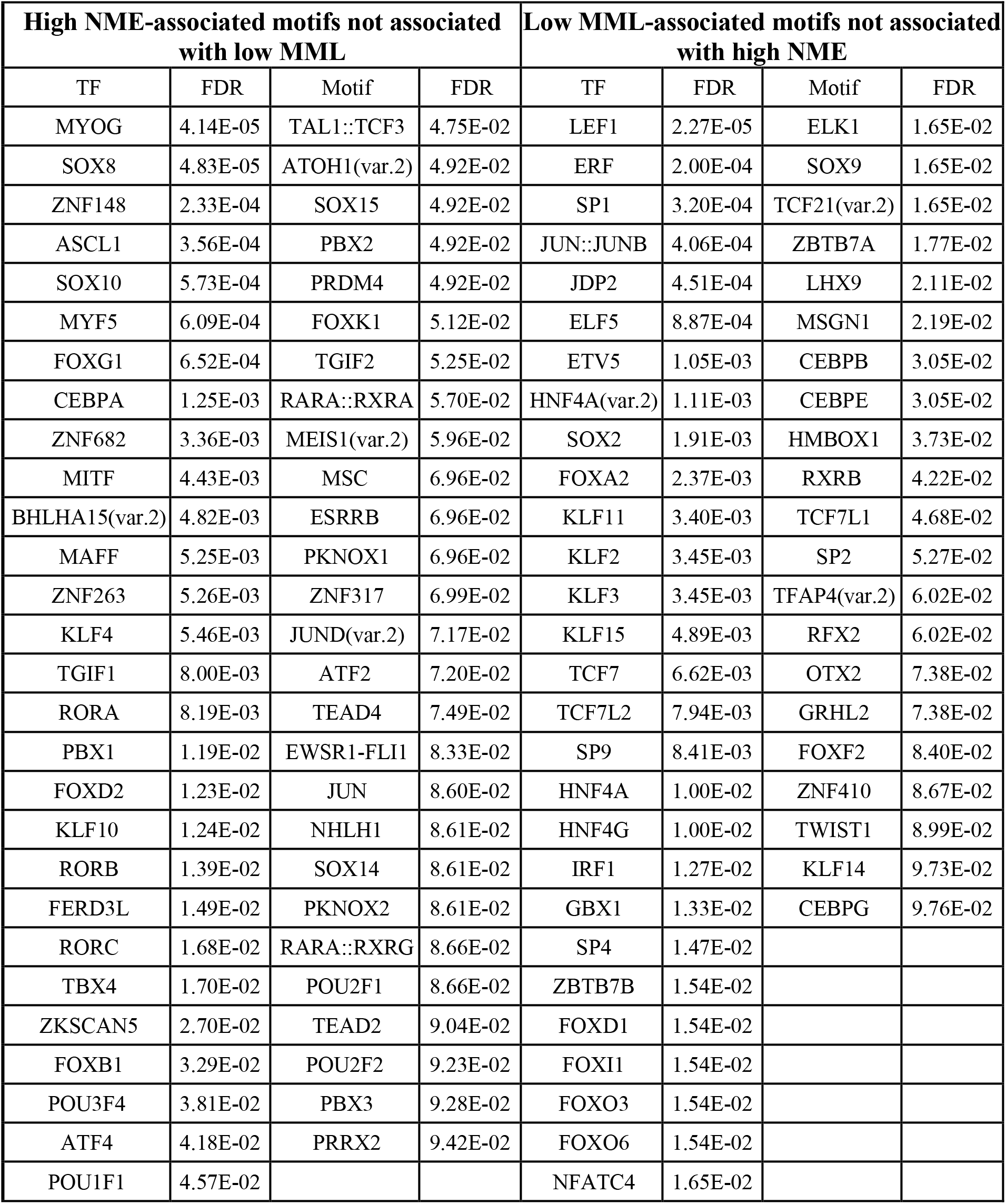
Transcription factors associated with high NME or low MML. The FDR is calculated from the P-values of the binomial test (P-values, 95% CI and effect size are in data S3). The GO enrichment analysis for the ranked list of transcription factors whose motifs only have high NME is *positive regulation of cell development* (enrichment=5.43, *P* =4.72 × 10^-5^) and *positive regulation of cell differentiation* (enrichment = 3.32, *P* = 7.71 × 10^-4^). Using the same GO enrichment analysis for the ranked list of transcription factors whose motifs only have low MML, there is no significantly enriched term using a P-value cutoff of 0.001.

Given that transcription factors generally bind to regulatory DNA to regulate their target genes (43), we then compared NME at regulatory DNA vs. non-regulatory DNA, defined using DNase I hypersensitive site sequencing data (see Methods and Materials). For 583 out of 630 motifs, the motif sites in regulatory DNA showed significantly higher NME than the motif sites in non-regulatory DNA (Wilcoxon’s signed-rank test, *FDR* < 0.05; Supplementary Figure S2A, B, and Supplementary Dataset S4). GO analysis of transcription factors for motifs with higher NME in regulatory DNA was also highly enriched for developmental categories (Supplementary Dataset S5). Examples include the LHX5 motif (Supplementary Figure S2C left panel), which regulates neuronal differentiation and dendritogenesis of Purkinje cells (44,45), and the SRF motif (Supplementary Figure S2C right panel), which is required for vascular smooth muscle cell differentiation (46). Low MML was also associated with regulatory DNA, as expected (Wilcoxon’s signed-rank test *FDR* < 0.05; and Supplementary Dataset S6 and Supplementary Figure S2D, E), e.g. CTCF (47) and NRF1 (48) (Supplementary Figure S2F).

### Entropy is inversely related to CpG density

We then performed allele-specific analysis to explore how normalized methylation entropy (NME) is related to DNA sequence other than transcription factor binding sites per se, by comparing allele-specific entropy to the trinucleotide DNA sequence contexts containing the given CpG dinucleotide (Figure 3A, Supplementary Figure S5; see Methods and Materials). This analysis revealed several such contexts, but the strongest effects were seen where the SNP results in a lost CpG in one allele, and the allele with fewer CpG was 2.1-fold more likely to show higher NME than the alternate allele (Figure 3A, Fisher’s exact test, 95% CI [2.0 - 2.2], *P* < 2.2 × 10^-16^). Comparing the number of CpG sites between the two alleles across all regions with significant allele-specific methylation entropy differences, the allele with more CpGs showed an average decrease in NME of 0.24 (Figure 3B, one-sided paired t-test, *P* < 2.2 × 10^-16^), supporting the observation that lower CpG number was associated with higher NME. We further examined NME across the genome within 20 kb of transcriptional start sites, regardless of allele, to include regions lacking SNPs. Here as well, there was an inverse correlation between NME and CpG density (Figure 3C top panel, Pearson correlation test, correlation = −0.21, 95% CI = [−0.21, −0.21], P < 2.2 × 10 ^-16^; see Methods and Materials). We also observed a negative correlation between the mean methylation level and CpG density (Figure 3C bottom panel, Pearson correlation test, correlation = −0.35, 95% CI = [−0.35, −0.35], *P* < 2.2 × 10^-16^). MML showed a more abrupt decrease at high CpG density than did NME (Figure 3C), consistent with the known compartmentalization of CpG islands.

### Information-theory based analysis reveals a developmental role of entropy

To explore the developmental significance of methylation information content, we analyzed mouse prenatal whole-genome bisulfite sequencing (WGBS) ENCODE3 data from 7 tissues at sequential embryonic developmental time points (E10.5 to E16.5) (10), examining 46 samples with data for 1035 pairwise sample comparisons at each of 5,017,785 genomic regions. We did not include postnatal time points since the environment of the animal is drastically different after birth, which could directly affect the epigenetic landscape, supported as well by the lack of a consistent developmental trajectory when a postnatal timepoint was included (Supplementary Figure S6). For each pairwise comparison, we calculated the information-theoretic distance between two samples’ methylation potential energy landscape, quantified by the uncertainty coefficient (UC). UC measures the inherent information in a sample distinct from another sample, corrected for entropy (mathmatically defined in the Methods).

Multidimensional scaling (MDS) of all 1035 comparisons across the entire genome showed that samples from the same tissue were grouped together, while samples from different tissues were separated (Figure 4A, top). Dissimilar tissue types, such as limb and liver, were separated from each other to a greater degree than similar tissue types, such as hindbrain, midbrain, and forebrain (Figure 4A top). Furthermore, MDS showed that samples from the same tissue type were ordered based on developmental stage, with samples from earlier stages being closer to a common center across tissues, and samples from later stages being away from the center, progressing in different directions for different tissue types (Figure 4A, bottom). Thus, the tissue-specific developmental trajectories can be derived from the information-theoretic content of DNA methylation.

**Figure 4.**
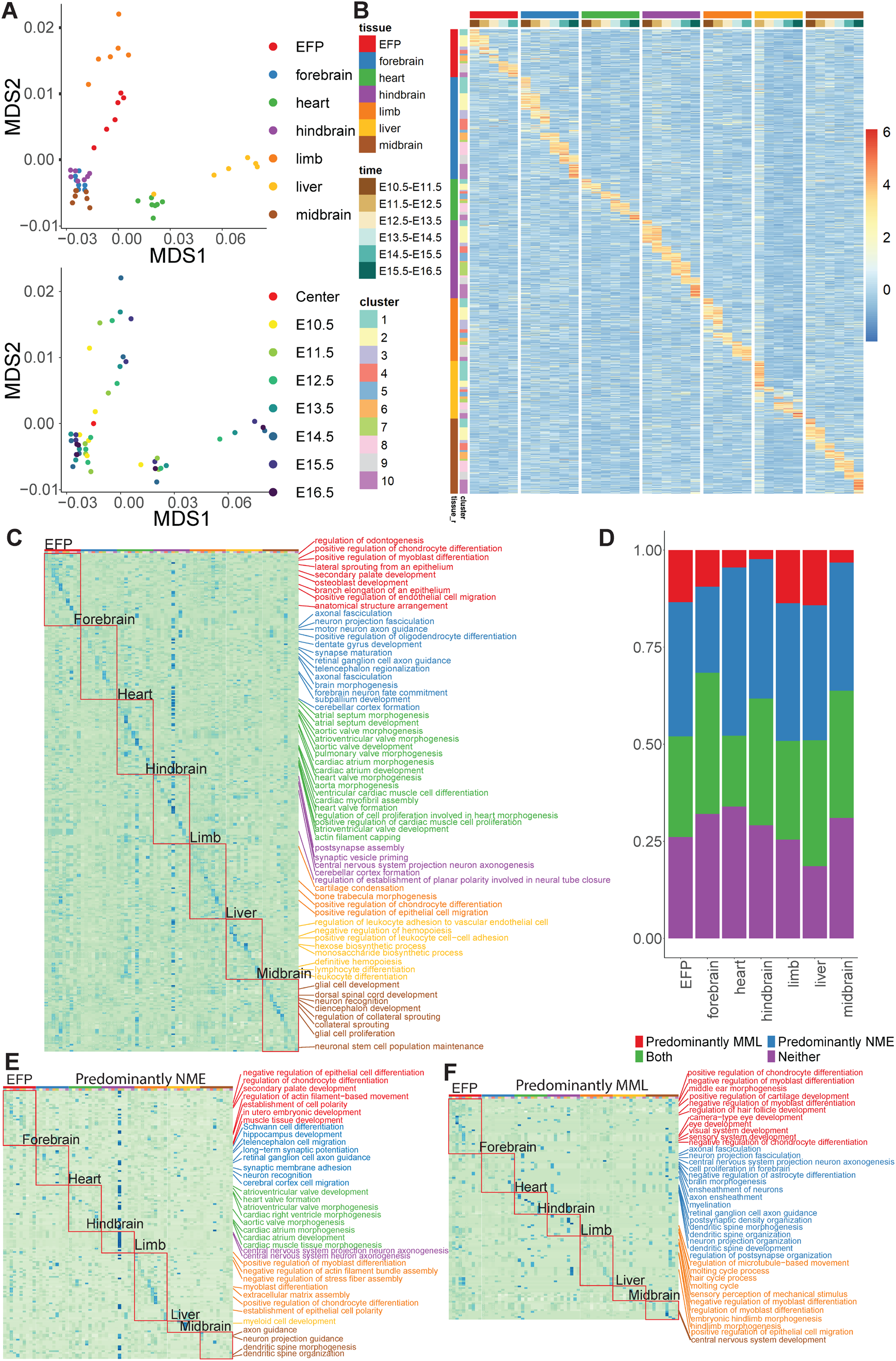
Information-theoretic analysis of methylation landscape in mouse embryonic development. **A.** Multi-dimensional scaling (MDS) plot of mouse embryonic samples from different tissues and developmental time points based on samples’ pairwise distance using uncertainty coefficient (UC). Samples (dots) are color-coded either by tissues (top) or by developmental time points (bottom). Samples from the same tissue or similar tissues tended to be clustered together. The MDS also captured samples’ temporal ordering, with samples from earlier developmental time points being closer to a common center (labeled by the red dot in the bottom plot) and samples from later time points gradually moving away from the center toward different directions representing different tissues. **B.** Heatmap of UC values in mouse embryogenesis revealed tissue- and developmental stage-specific DNA methylation landscape changes. Regions with high UC in only one tissue were grouped into 10 clusters, which were ordered according to the consecutive time points with the highest UC. The clustering reflected the temporal cascade of methylation landscape changes in that tissue. High UC (red) means large methylation change (including mean and entropy changes) and low UC (blue) means small change. The temporal patterns were tissue-specific, as patterns observed in one tissue cannot be observed in other tissues. **C.** Gene ontology (GO) analysis demonstrates the biological significance of UC-based developmental clustering. For each tissue and region cluster identified in (B), enriched biological functions associated with enhancer regions in the cluster are identified and the top 5 GO terms with largest fold-enrichment are shown (rows: GO terms; columns: tissues and region clusters; “*”: FDR<=0.1). Large fold-enrichments were labeled in blue while small fold-enrichments were labeled in green. The enriched GO terms revealed tissue-specific functions. (Note: although GO terms in cluster 6, hindbrain have high fold-enrichments in other tissues, they are not significant.) The GO terms in the heart followed the development order, i.e. we observed atrial septum and cardiac atrium development in early stages and heart valve development in late stages. Similarly, in the forebrain, we saw the GO terms related to neuron development in early stages and brain subdivision formation in later stages. **D.** Bar plot showing the relative contribution of MML and NME to the methylation landscape changes as characterized by UC. In each tissue, regions were classified into four categories based on whether a region’s UC change across time points was predominantly correlated with the change in MML (red), NME (blue), both (green), or uncertain (purple). The proportion of each category among all regions that have high UC in at least one pair of time points is shown. In all tissues, there were more regions where UC is predominantly correlated with NME change (blue) than the regions where UC is predominantly correlated with MML change (red), suggesting a larger role of methylation entropy than mean methylation in shaping the dynamic changes of methylation landscape. **E.** Gene ontology (GO) analysis demonstrates the biological significance of regions whose UC is predominantly correlated with NME using the same method as in **C**. All tissues show tissue-specific GO terms from those regions whose UC is predominantly correlated with NME. **F**. Gene ontology (GO) analysis demonstrates the biological significance of regions whose UC is predominantly correlated with MML using the same method as in **C**. Some tissues show tissue-specific GO terms such as EFP, forebrain, limb and midbrain. In contrast to **E**, some tissues do not show any tissue-specific GO terms, including heart, hindbrain, and liver.

We then performed clustering analysis within each tissue in order to determine whether there was a specific set of genomic regions undergoing methylation landscape changes at each developmental stage, and if so, whether those time-specific clusters were also specific to the given tissue in which they were observed. To do this, for each tissue, we performed clustering analysis on all pairs of adjacent time points from earliest to latest, analyzing 2,085,884 regions showing large methylation landscape change (defined by UC > 0.1 in at least one tissue between any two time points, see Methods and Materials, Supplementary Figure S7) for each comparison. The ordering of the regions in the clustering heatmap recaptured the developmental process, confirming that regions with high UC capture developmental change (Supplementary Figure S7). The temporal patterns were tissue-specific, as clustering patterns observed in one tissue were not observed in other tissues. To explore the background signal in Supplementary Figure S7, we also measured the proportion of regions with high UC differences between any time point pair, by the number of tissues in which it was observed, and this was bimodal with the highest peak for unique tissues (Supplementary Figure S8). These tissue-specific regions with high UC also captured time-dependent developmental changes (Figure 4B). Furthermore, these developmental clusters in a given tissue were almost entirely specific to that tissue (Figure 4B). UC > 0.1 corresponded to the top 5% of all UC values in the entire dataset. Using other UC cutoffs from 0.025 to 0.2, we still observed the tissuespecific clustering patterns. (Supplementary Figure S9, see Methods and Materials).

Next, we asked whether the genes linked to these developmental- and tissue-specific regions are functionally related to those tissues by GO analysis. We mapped the regions to regulatory elements of specific genes, including promoters (within 2 kb of the transcriptional start site), and enhancers as described (24). Enhancers showed substantially larger changes in both NME and MML than promoters (Supplementary Figure S10), so we focused initially on enhancer regions (Figure 4C, Supplementary Figure S11 and Supplementary Dataset S7). The GO terms identified in this manner were strikingly tissue-specific as well as specific to each temporal cluster, and highly related to the development function of each tissue as well. For example, GO terms related to embryonic facial prominence (EFP), such as positive regulation of chondrocyte differentiation, showed large and significant enrichment only in EFP and not in other tissues; and GO terms related to liver, such as monosaccharide biosynthetic process, showed large and significant enrichment only in liver and not in other tissues (Figure 4C). Moreover, in some tissues, the temporal order of GO terms also reflected the temporal developmental program (forebrain and heart, see Figure 4C). Using the same method with other UC cutoffs from 0.025 to 0.2, we still found significant tissue- and time-specific GO enrichment (Supplementary Figure S12).

Although our focus was on enhancer regions, as explained above, we also analyzed the genes whose promoters showed significant methylation landscape change. We found that those genes were enriched in fewer tissue-specific terms than genes with methylation landscape changes in enhancer regions (Supplementary Figure S13). This was consistent with the finding that enhancers showed larger NME and MML changes than promoters (Supplementary Figure S10), confirming that epigenetic landscape changes at enhancers were more strongly related to tissue-specific development.

### UC is more strongly correlated with entropy than with mean methylation

To determine the relative contribution of mean methylation and methylation entropy change to the developmental landscape distance as measured by UC as described above, we next calculated the correlation between MML change and NME change to UC, classifying methylation changes at each region as predominantly correlated with MML, NME, both, or neither (see Methods and Materials). In the regions with UC > 0.1 between developmental timepoints, UC was predominantly correlated with MML change in 2%-14%, depending on the tissue, with NME change in 22-43%, with both MML and NME change in 18-36% and with neither in 19%-34% regions (Figure 4D). These data suggested that in all seven tissues, there was a much larger contribution of methylation entropy changes compared to methylation mean changes to the dynamic methylation landscape in development (one-sided t-test, 95% CI = [14659, Inf], *P* = 1.8 × 10^-6^). Even using other UC cutoffs from 0.025 to 0.2, we still observed a larger contribution of methylation entropy than methylation mean to the dynamic methylation landscape in development (Supplementary Figure S14).

To understand the biological function of those regions with different categories as shown in Figure 4D, we performed a GO annotation analysis on the genes with enhancers in the four UC-correlation categories described above. Predominantly NME-correlated regions were overall more enriched for tissue-specific functions than predominantly MML-correlated regions (Figure 4E, F, Supplementary Figure S15A, B and Supplementary Dataset S8-9), suggesting the NME and MML change are related to different functions in mouse embryonic development and the NME change showed greater correlation with UC than MML change. There was also substantial enrichment for tissue-specific functions in UC regions correlated with both mean and entropy change (Supplementary Figure S15C, and Supplementary Dataset S10), and UC also captured some information independent of both mean and entropy change (Supplementary Figure S15D, and Supplementary Dataset S11). These results demonstrated that methylation entropy contains at least as much information about the developmental epigenetic landscape as does mean methylation. Thus, analyzing only mean methylation prevents one from identifying the majority of DNA methylation landscape changes characterized by the information-theoretic measure UC.

### Developmental entropy and mean are associated with different transcription factor binding sites

Given the results above showing that mean and entropy were associated with different transcription factor motifs in human, and also associated with different genomic regions based on embryonic development, we asked whether different transcription factor motifs are associated with mean- or entropy-related developmental methylation changes in the mouse. We compared regions where UC was predominantly correlated with NME change, to regions where UC was predominantly correlated with MML change (Figure 5, see Methods and Materials). Different sets of motifs were found to be associated with entropy versus mean. In total, we identified 344 entropy-associated motifs and 324 mean-associated motifs (Supplementary Dataset S12). Figure 5 shows the enrichment of each motif in the regions where UC was predominantly correlated with NME change versus the regions where UC was predominantly correlated with MML change in all 7 tissues we analyzed. For example, KLF family motifs such as KLF4 and KLF5 are among the entropy-associated motifs which appear in multiple tissue types such as heart, forebrain, limb, EFP, midbrain, hindbrain, and liver (Figure 5A-G, Supplementary Dataset S12). KLF4 is known to be a key transcription factor in embryonic development and is also one of the four transcription factors to generate iPSC (49). Among mean-associated motifs, NF-I family motifs such as NFIC, NFIX, and NFIB were identified in multiple tissues such as heart, limb, EFP, hindbrain, and liver (Figure 5A, C, D, E, F, and Supplementary Dataset S12). Mutations in *Nfi* family members are associated with brain development defects (50). Using entropy-associated motifs, we annotated enhancers with entropy changes as well as their target genes (Table 2, Supplementary Dataset S13, and Supplementary Figure S16), and similarly for enhancers with mean methylation changes (Supplementary Dataset S14). For instance, *Celsr2*, which has been shown to regulate forebrain wiring (51), showed a large entropy change between E12.5 and E13.5 at its enhancer, which is also the KLF4 transcription factor binding site (Supplementary Figure S16) (49). Similarly, *Gata5* plays a critical role in heart development and *Gata5* knockout mice develop bicuspid aortic valve (52). *Gata5* showed a large entropy change between E11.5 and E12.5 at its enhancer, which is also the GLI3 binding site (Supplementary Figure S16) (49). These data suggest that entropy-associated transcription factor binding may play an important role in modulating enhancer function in development.

**Figure 5.**
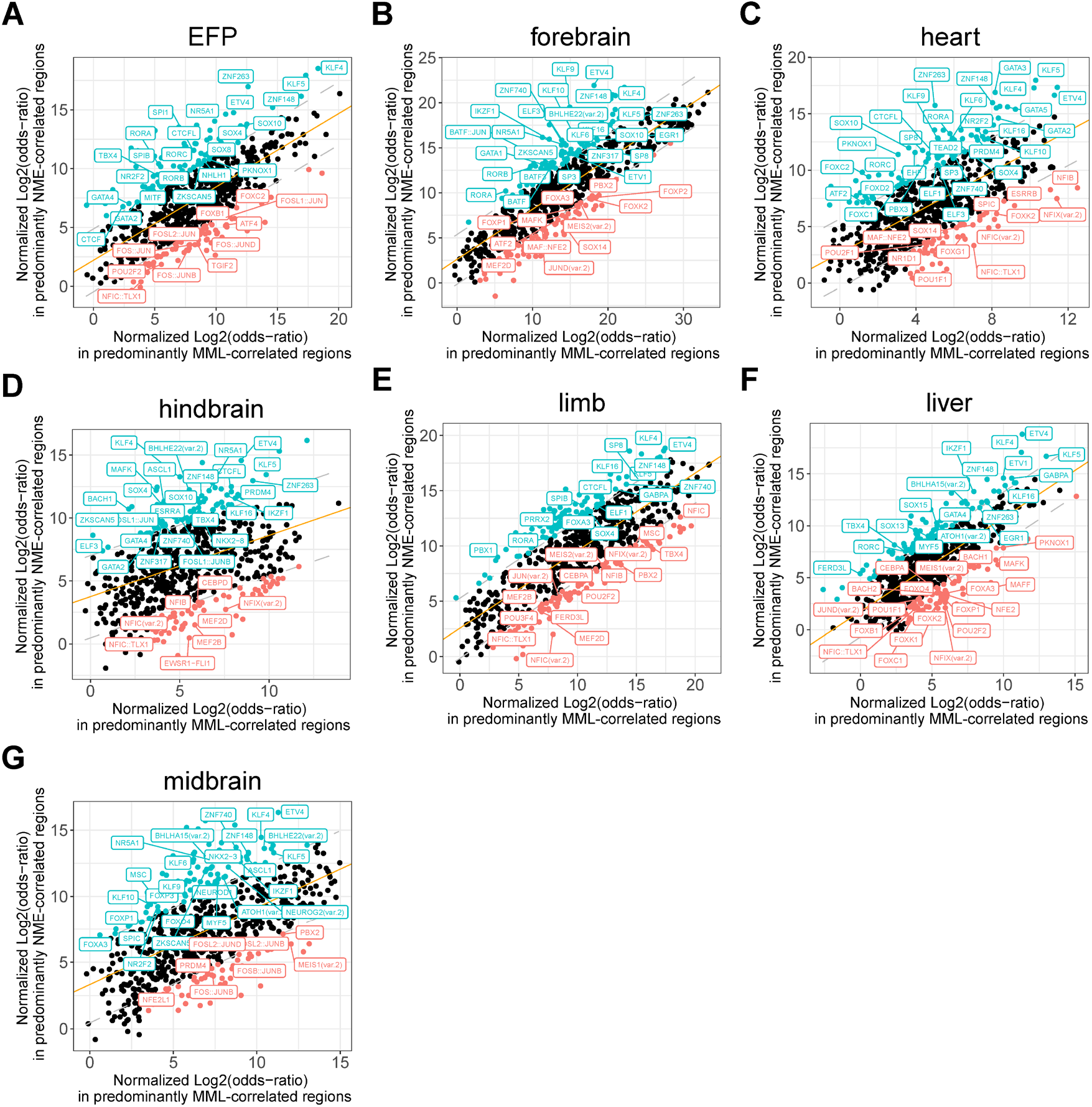
Association of methylation entropy changes with transcription factor binding motifs. Scatterplots comparing the enrichment level of each TF motif in regions where DNA methylation changes were predominantly correlated with NME change versus the enrichment level of the same motif in regions where methylation changes were predominantly correlated with MML change. The analysis was run for **A.** embryonic facial prominence (EFP), **B.** forebrain, **C.** heart, **D.** hindbrain, **E.** limb, **F.** liver, **G.** midbrain, in the mouse embryonic development dataset. For each tissue, we identified motifs that were more enriched in the regions where UC was predominantly correlated with NME change compared to regions where UC was predominantly correlated with MML change (entropy-associated motifs, marked with blue color) as well as motifs that were more enriched in regions where UC was predominantly correlated with MML change than the regions where UC was predominantly correlated with NME change (mean-associated motifs, marked with red color). Motifs that were also identified in the human analysis which prefer high NME are marked with text labels. Entropy-associated motifs that prefer high NME in the human analysis are marked with blue labels. Mean-associated motifs that preferred high NME in the human analysis are marked with red labels. Among the entropy-associated motifs, a substantial number (32, 27, 16, 24, 26, 27, and 19 motifs from heart, forebrain, limb, EFP, midbrain, hindbrain, and liver, respectively) also prefer high NME in the human analysis such as KLF4 and KLF5.

**Table 2.**
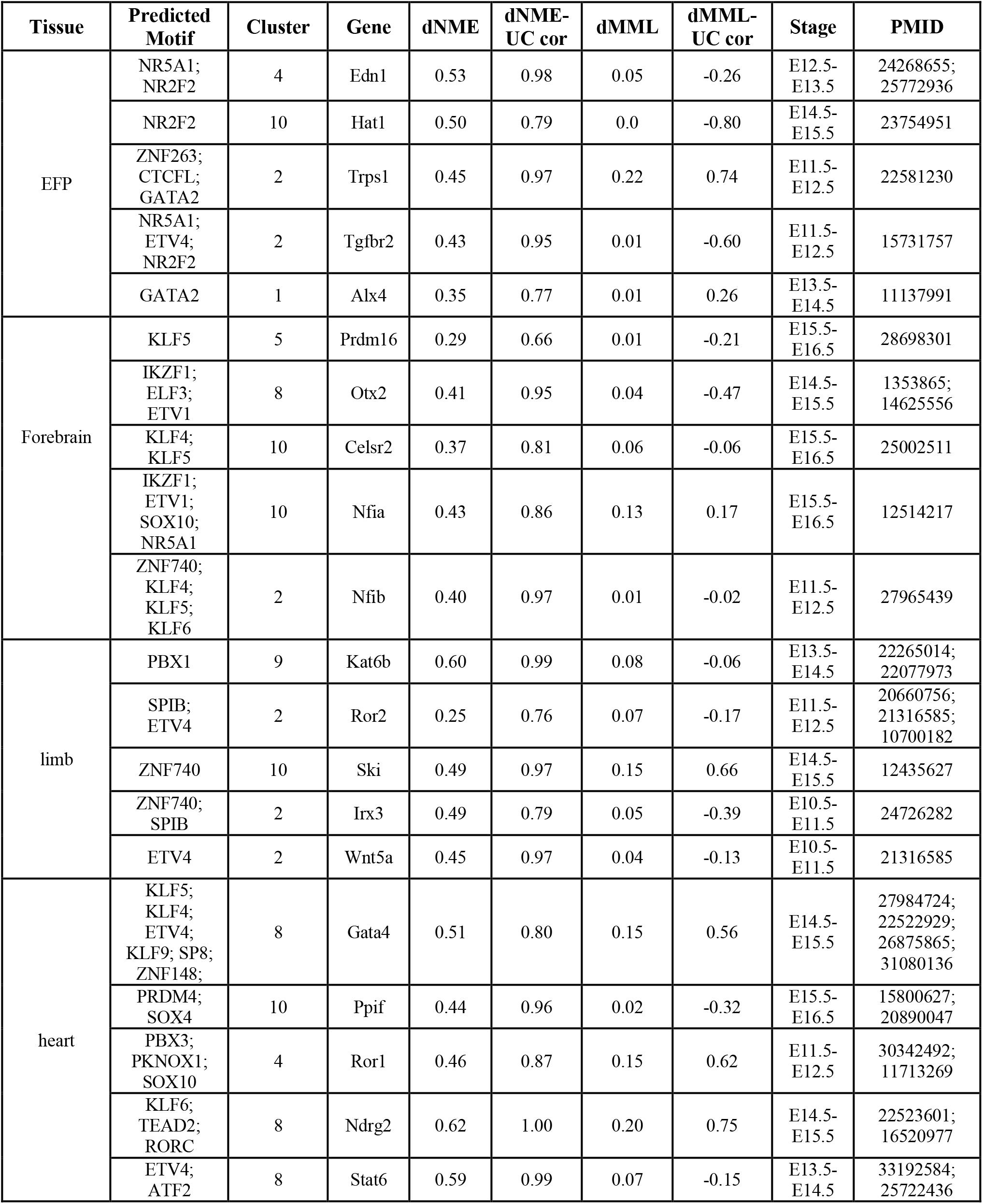
Genes with enhancers contain entropy-associated transcription factor motifs. Genes whose enhancers contain entropy-associated motifs, are listed in the “Entropy-associated TF motif” column. At the enhancer of those genes, the correlations between UC and NME difference (dNME) are much greater than the correlation between UC and MML difference (dMML) and the dNME is much larger than dMML. Those genes are known to be functional in the corresponding tissue from the literature listed in the PMID column. This suggests that those entropy-associated motifs are at the enhancers of functional genes.

### Entropy-associated sequence and its association with regulatory DNA are conserved between human and mouse

Next, we asked whether the entropy-associated sequence and chromatin features are conserved between human and mouse. The percentage of overlap between human motifs associated with high NME and mouse entropy-associated motifs was significantly higher than the percentage of overlap between the human motifs associated with high NME and the mouse MML-related motifs (Fisher’s exact test, odds ratio = 1.43, *P* = 0.04; Supplementary Dataset S12). The mouse entropy-associated motifs that overlap with the human motifs preferring high NME are highlighted in blue boxes in Figure 5. Those entropy-associated motifs can play a critical role in the corresponding tissue, such as PBX3, whose entropy-associated binding motifs are conserved in human and mouse, and are associated with heart development but not forebrain and limb, is necessary for normal heart development (53), PKNOX1, which is associated with congenital heart defects (54) and FOXC1, which is required for cardiovascular development (55). In the forebrain, we found SOX10, SP8, and RORA (Figure 5B), which regulates myelination-related genes (56), promotes olfactory bulb interneurons during the development (57), and regulates genes associated with autism spectrum disorder (58). In limb, we found SOX4, which is related to massive cartilage fusion (59), ETV4, associated with sonic hedgehog pathways in limb outgrowth (60), and PBX1, required for chondrocyte proliferation (61).

We found similar conservation between mouse and human for the relationship of entropy to CpG density and regulatory DNA. For CpG density, similar to human (Figure 3C), we observed a strong correlation between local CpG density and NME in mouse samples (Pearson correlation test, correlation = −0.21, 95% CI = [-0.21, −0.21], *P* < 2.2 × 10^-16^; Supplementary Figure S17). To explore the relationship between NME and transcription factor motif binding sites at regulatory DNA in mouse, we compared the NME between motif sites that are in regulatory DNA as defined by a union set of mouse DNase I hypersensitive sites and those located in non-regulatory DNA in each mouse sample for 736 TF motifs. Similar to human motif analysis (Supplementary Figure S2), we found that most of the motifs in mouse have significantly higher NME in regulatory DNA than non-regulatory DNA (661 motifs with *FDR* ≤ 0.05) (Supplementary Figure S18, and Supplementary Dataset S15).

Together, these results showed that the relationships between NME and transcription factor binding, CpG density, and regulatory DNA were highly conserved between mouse and human, thus supporting their functional significance.

### Methylation entropy is associated with gene expression variability in both human and mouse

Because NME is associated with expression variability in human cancer cells (62), we asked whether methylation entropy is related to gene expression variability in normal human somatic tissues and mouse embryonic development. First, we re-analyzed singlecell RNA-seq data (26) from 14 different human somatic tissues and human ESC that we had already done methylation analysis from WGBS data for the corresponding tissues (see Materials and Methods for details). We found a correlation between NME near the transcription start site (TSS) of the genes and mean-adjusted variability (MAV) of gene expression (see Methods and Materials). Genes with higher MAV tended to have larger NME near TSS while genes with lower MAV tended to have smaller NME near TSS (Figure 6A) (26). The correlation between the NME close to the TSS and the gene expression variability was highly significant (Pearson correlation test, average correlation = 0.23, average 95% CI = [0.21, 0.25], average *P* = 7.6 × 10^-5^). We observed a similar pattern in other human samples except for undifferentiated stem cell lines (Figure 6B). Thus, NME was generally positively correlated with gene expression variability but not mean expression (Supplementary Figure S19).

**Figure 6.**
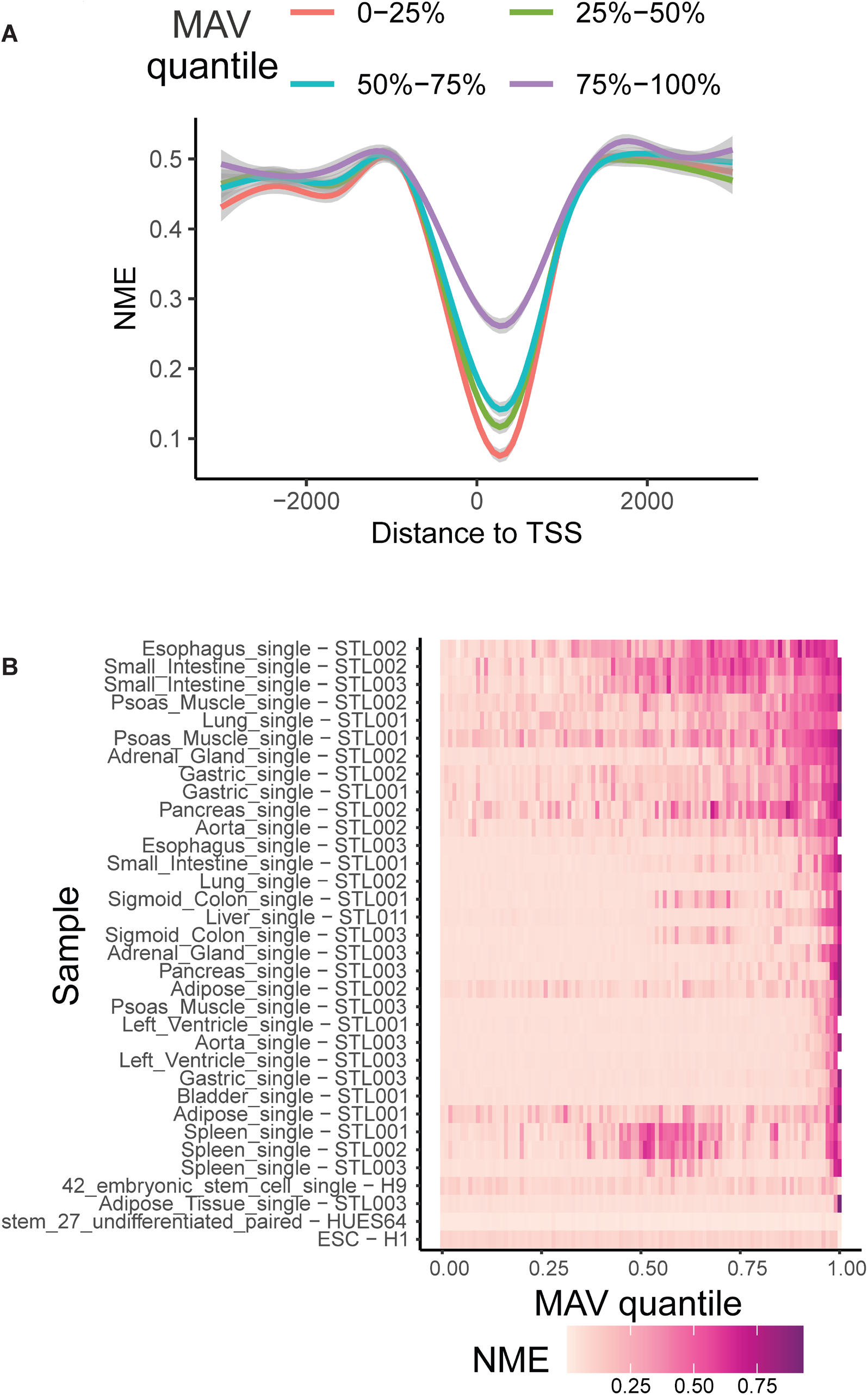
Association of methylation entropy changes with gene expression variability. **A.** Association between NME surrounding transcription start sites (TSS) and gene expression variability. An example showing NME near TSS was lower for the gene with low mean-adjusted expression variability (MAV) than the gene with higher MAV. Genes were stratified based on the quartiles of their MAV. For each stratum, the average NME surrounding TSS across all genes in the stratum is shown. The genes in the lowest MAV quantile (red line) had the lowest NME near their TSS. The genes in the highest MAV quantile (purple line) had much higher NME near their TSS. The Pearson correlation between genes’ MAV and NME was 0.269 (P < 2.2 × 10^-16^). **B.** Heatmap showing NME values within +-500 bp of genes’ TSS in different samples. Each row is a sample. In each sample, genes were stratified based on their expression variability using quantiles of MAV (X-axis), and the heatmap shows the average NME of all genes in each stratum. We observed increasing NME (from pink to dark red) as the MAV quantile increases except for embryonic cell lines, indicating that gene expression variability (MAV) is correlated with NME in regions surrounding genes’ TSS.

Given the enrichment of regions with large MML difference in imprinted genes, we also explored the relationship between MML and monoallelic gene expression (MAE)(18). We observed significant enrichment of the regions with large allelic MML difference in the promoters of MAE genes (Fisher’s exact test, odds ratio = 1.49, 95% CI=[1.04 – 2.15], *P* = 0.03, Supplementary Dataset S16). In contrast, as expected, we did not observe an enrichment of the regions with large allelic NME difference (Fisher’s exact test, odds ratio = 1.03, 95% CI = [0.79 – 1.35], *P* = 0.84, Supplementary Dataset S16) suggesting no association between methylation entropy and mean gene expression. Thus, while mean methylation is associated with mean expression, entropic methylation is associated with expression variability (MAV).

We next analyzed the relationship between NME and MAV in mouse, analyzing mouse fetal limb tissue scRNA-seq data from the ENCODE3 project (28) and calculating MAV in the same way as in human. Consistent with the human analysis, genes with higher MAV showed larger NME near TSS while genes with lower MAV showed smaller NME near TSS (Supplementary Figure S20A), and there was a smaller but still statistically significant correlation between NME and MAV (Pearson correlation test, average correlation=0.11, average 95% CI = [0.10, 0.11], average *P* < 2.2 × 10^-16^; Supplementary Figure S20B) (28). Those observations showed that the association of NME and gene expression variability in normal tissue is also conserved between human and mouse.

## Discussion

In summary, we have performed a systematic characterization of DNA methylation entropy, relating it to DNA sequence, predicted transcription factor binding sites, regulatory DNA, its relative contribution to epigenetic landscape changes during mouse development, and its relationship to variable gene expression. Applying an information-theoretic approach to mouse embryo DNA methylation data, we identified tissue- and time-specific transitions that are much more discriminative than conventional mean-based DNA methylation analyses, identifying temporal transitions remarkably well and essentially uniquely for each tissue type. By using the information metric of UC, we could also directly compare the relative contribution of differences in entropy versus mean methylation to these informational changes, and entropy changes were 2- to 3-fold more abundant than mean differences in the epigenetic landscape of mouse development. Gene ontology enrichment analysis supported a functional role of these entropy changes, particularly at enhancers, identifying many known critical genes for development in those tissues. These specific functional categories were often congruent to the functional development of those tissues, as well, as evidenced particularly in the heart.

We also identified certain DNA sequence relationships with methylation entropy, using allele-specific analysis comparing directly the entropy or mean of sequences differing by a single nucleotide in the same cells. There were approximately 3-fold more entropy- than mean-distinguishing SNPs. Moreover, these sequence differences were directly related to changes in many transcription factor binding sites. Note that imprinted genes were associated with differential mean methylation between alleles, although if the alleles are pooled they will appear entropic as in (11). There was also a strong inverse relationship of entropy with small differences in CpG density, in contrast to the comparatively weak inverse association of CpG density with mean methylation except at very high-density islands.

Moreover, entropic methylation was strongly associated with gene expression variability in human normal somatic cells and normal mouse development, in contrast to the monoallelic expression which was associated with differential mean methylation as seen here and elsewhere (11,63). The fact that so many entropy-associated features were found to be conserved between human and mouse suggests they serve an important functional role. This relationship supports a potential role of epigenetic entropy in mediating either buffering or phenotypic divergence, two sides of the same developmental coin (4). The link between epigenetic entropy and the sequence of specific transcription factor binding sites suggests that binding of some transcription factors may regulate a variable response to environmental signaling, since many of these transcription factors are signaling-dependent, as classified by Brivanlou and Darnell (64). These include nuclear receptor family members such as NR5A1, RORA, RORB, and accessory factors like KLF’s, and receptor-ligand targets including ETS-family members (ETV’s, ELF’s), receptor tyrosine kinases like EGFRs, and HIPPO-signaling targets like TEADs, all of which were conserved entropy-associated binding regions between human and mouse.

A limitation of the mouse data analysis is inherent in the datasets from which they were derived (and as those authors also acknowledged as a limitation) (10), namely that a given tissue sample would include multiple cell types which could affect inferences about developmental changes. This admixture would likely affect conclusions regarding entropy changes over time in development. However, entropy itself likely does contribute meaningfully to developmental transitions based on the current analysis for two reasons. First, the regions associated with high entropy change are located at the enhancers of the genes that are specific to the development of corresponding tissue as shown in the gene ontology (GO) analysis. Second, this limitation of admixture does not apply to the human data sets since in that case entropy differences were at the level of individual alleles. Moreover, the entropy-associated transcription factor motifs were conserved between the human data set and the mouse embryonic data set.

It will be interesting to explore the relationship between polymorphisms and epigenetic entropy in large cohorts, which could not be done on the relatively small number of human samples that we were able to analyze here, since rare variants in transcription factor binding sites have been associated with altered methylation at specific CpG sites in human lymphocytes (65). Previously, Mendelian transmission of 383 regions with variable mean methylation in the human genome were identified (66), and two recent studies using GTEx data show SNPs associated with variation in mean methylation in a tissue-specific manner (67,68). An alternative approach would be to analyze genetically complex mouse strains like the collaborative cross (69,70), which was not possible in the inbred mouse strain studied here; and this mouse approach would allow for hypothesis testing of specific transcription factor engineered mutations in the development.

Finally, the results shown here emphasize the importance of including entropy analysis in genome-wide studies of DNA methylation. While this has been widely adopted in some human diseases such as cancer (71), the role of methylation entropy in embryonic development may be under-appreciated. A recent study supports that idea, showing that knockout of DNA methyltransferases increases methylation entropy and gene expression variability in cultured embryonic stem cells (7). The results presented here identify genetic sequences and their associated motifs underlying epigenetic entropy and their critical role in development.

## Data and materials availability

All the code for analysis and data are available through the data file submitted along with the manuscript and the AWS S3 bucket, which anonymizes access: s3://methlandscape/ using AWS Command Line Interface (CLI) with the following credentials: Access key: AKIAYIZXZQF2XNNPRWE3 and private access key for reviewers: dnhJwHvpdnLvdkBa0AlFXbcTfRB3/Mdp7NG8Obdx. Mouse data are from the ENCODE3 project: https://www.encodeproject.org/. Human data are from the ENCODE project and Roadmap Epigenetic project: http://www.roadmapepigenomics.org/

## Acknowledgments

We thank Mauro Maggioni, Bill Nelson, Sarah Wheelan, Patrick Cahan, and Nilanjan Charterjee for their careful review and thoughtful suggestions.

