## Supplementary Figures for "DNA methylation entropy is associated with DNA sequence features and developmental epigenetic divergence"

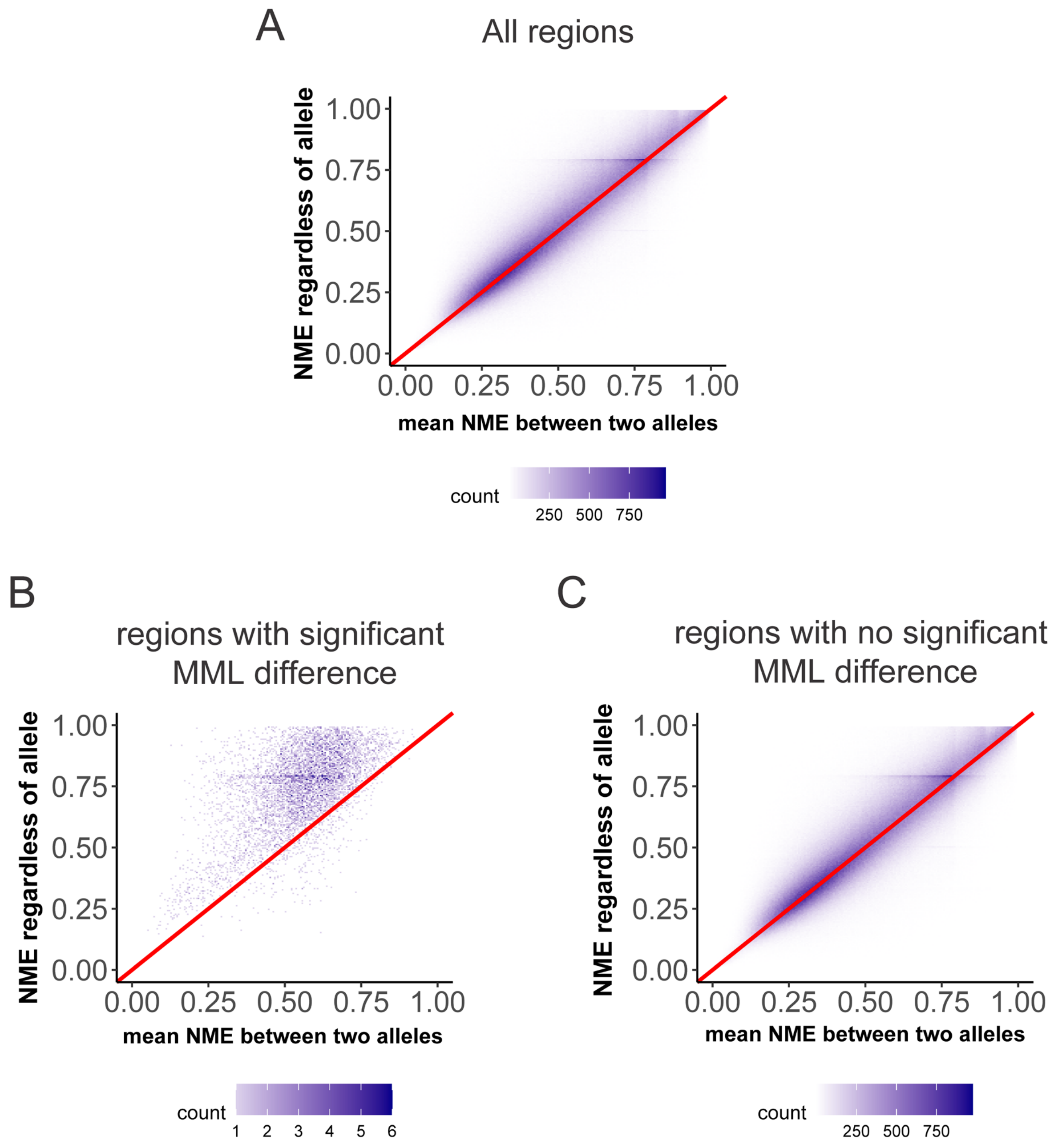


**Fig. S1. Relationship between NME regardless of allele and mean allelic NME. A**. Dot plot showing that the mean allelic NME and NME regardless of the allele are highly correlated across all genomic regions. The red line is the diagonal line. Most of the dots are very close to the diagonal line. **B**. The dot plot shows that NME, regardless of allele, is higher than mean allelic NME at regions that show a strong mean methylation level difference between two alleles. **C.** The dot plot shows that NME, regardless of allele, and mean allelic NME are highly correlated in the regions that do not show strong mean methylation differences between the two alleles.

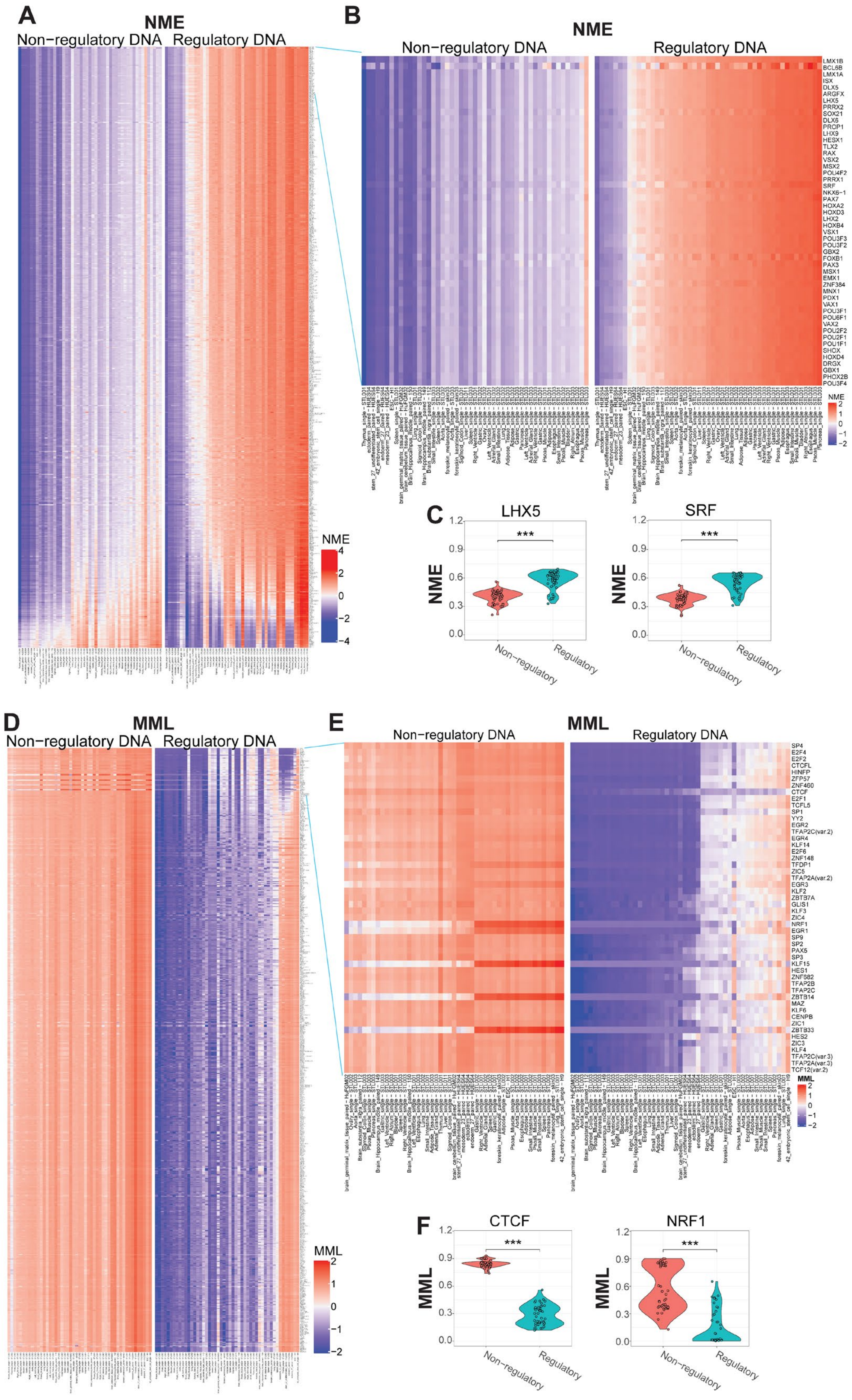

A
NME
Non−regulatory DNA
Regulatory DNA
NME
4
2
0
−2
−4
B
NME
Non−regulatory DNA
Regulatory DNA
LMX1B
BCL6B
LMX1A
ISX
DLX5
ARGFX
LHX5
PRRX2
SOX21
DLX6
PROP1
LHX9
HESX1
TLX2
RAX
VSX2
MSX2
POU4F2
PRRX1
SRF
NKX6−1
PAX7
HOXA2
HOXD3
LHX2
HOXB4
VSX1
POU3F3
POU3F2
GBX2
FOXB1
PAX3
MSX1
EMX1
ZNF384
MNX1
PDX1
VAX1
POU3F1
POU6F1
VAX2
POU2F2
POU2F1
POU1F1
SHOX
HOXD4
DRGX
GBX1
PHOX2B
POU3F4
NME
2
1
0
−1
−2
C
LHX5
SRF
NME
***
1.2
0.9
0.6
0.3
0.0
Non−regulatory
Regulatory
D
MML
Non−regulatory DNA
Regulatory DNA
MML
2
1
0
−1
−2
E
MML
Non−regulatory DNA
Regulatory DNA
SP4
E2F4
E2F2
CTCFL
HINFP
ZFP57
ZNF460
CTCF
E2F1
TCFL5
SP1
YY2
EGR2
TFAP2C(var.2)
EGR4
KLF14
E2F6
ZNF148
TFDP1
ZIC5
TFAP2A(var.2)
EGR3
KLF2
ZBTB7A
GLIS1
KLF3
ZIC4
NRF1
EGR1
SP9
SP2
PAX5
SP3
KLF15
HES1
ZNF582
TFAP2B
TFAP2C
ZBTB14
MAZ
KLF6
CENPB
ZIC1
ZBTB33
HES2
ZIC3
KLF4
TFAP2C(var.3)
TFAP2A(var.3)
TCF12(var.2)
MML
2
1
0
−1
−2
F
CTCF
NRF1
MML
***
1.2
0.9
0.6
0.3
0.0
Non−regulatory
Regulatory

**Fig. S2. Transcription factor (TF) motif sites have higher normalized methylation entropy (NME) and lower mean methylation level (MML) at regulatory DNA than non-regulatory DNA. A.** Heatmaps comparing NME between regulatory DNA (right panel) and non-regulatory DNA (left panel) for the 630 motifs from the JASPAR database. Each row corresponds to a TF binding motif and each column corresponds to a WGBS sample. For each motif and in each WGBS sample, the median NME across all regulatory (right panel) or non-regulatory (left panel) DNA with the motif is shown. For visualization, NME values within each row were standardized across regulatory and non-regulatory DNA to have zero mean and unit SD (red, high NME; blue, low NME). **B.** Zoomed in heatmaps comparing NME between regulatory DNA (right panel) and non-regulatory DNA (left panel) for the top 50 motifs in **A** with the largest difference between regulatory and non-regulatory DNA. **C.** Violin plots comparing NME between regulatory and non-regulatory DNA for two example motifs (LHX5 and SRF). For both motifs, the NME of regulatory motif sites was significantly larger than the NME of the non-regulatory motif sites (Wilcoxon's signed-rank test FDR < 0.05 for both LHX5 and SRF). **D.** Heatmaps comparing MML between regulatory DNA (right panel) and non-regulatory DNA (left panel) as in **A** (red, high MML; blue, low MML). **E.** Zoomed in heatmaps comparing MML between regulatory DNA (right panel) and non-regulatory DNA (left panel) for the top 50 motifs with the largest difference between non-regulatory and regulatory DNA as in **D**. **F.** Violin plots comparing MML between regulatory and non-regulatory DNA for two example motifs (CTCF and NRF1). MML of regulatory motif sites was significantly larger than MML of non-regulatory motif sites (Wilcoxon's signed-rank test FDR < 0.05) for both CTCF and NRF1.

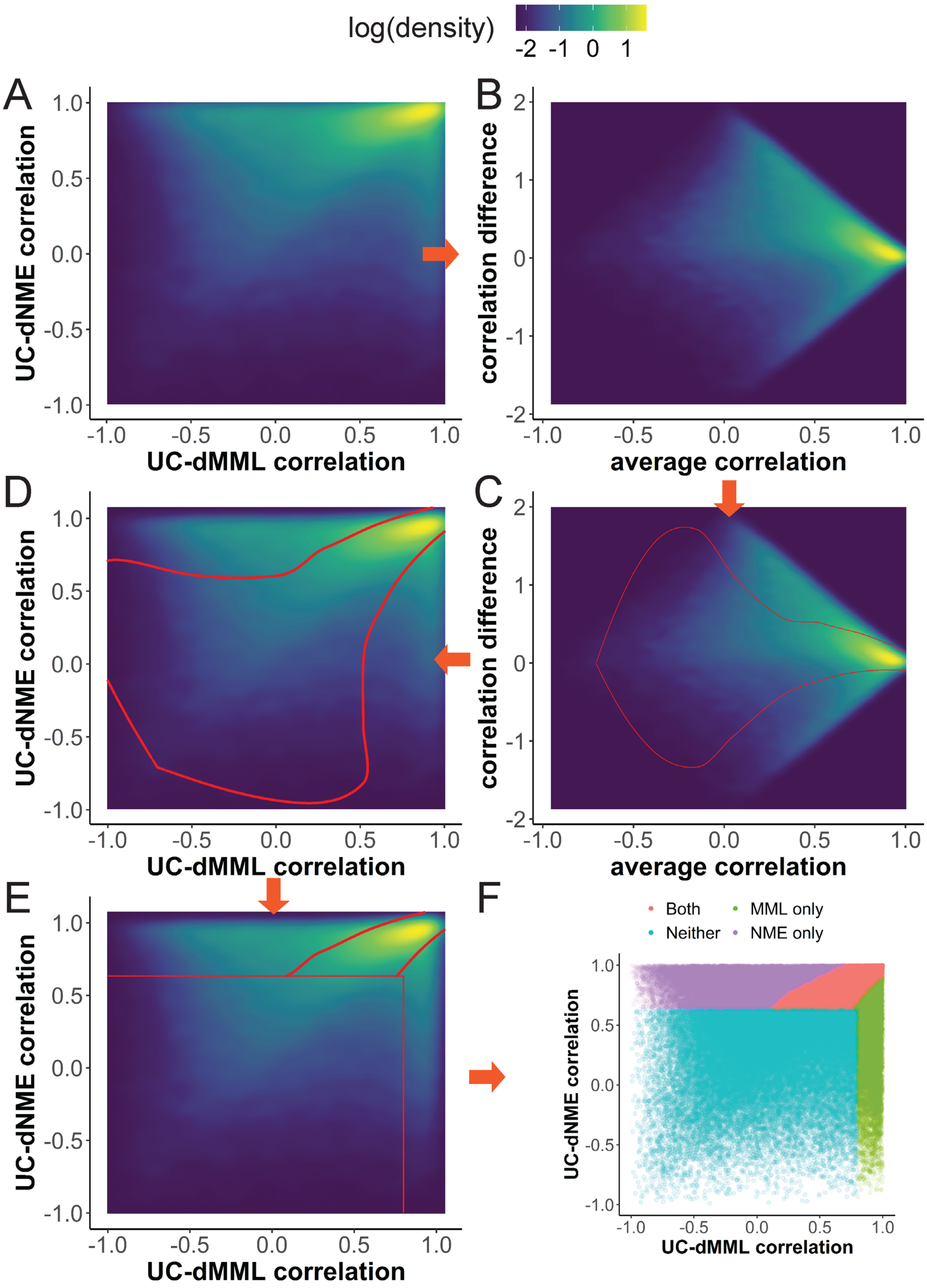

log(density)
-2 -1 0 1
A
B
C
D
E
F
UC-dNME correlation
UC-dMML correlation
correlation difference
average correlation
Both
MML only
Neither
NME only

**Fig. S3. Categorizing regions in the mouse. A.** Density plot shows the relationship between dMML-UC correlation and dNME-UC correlation. **B.** difference-average (M-A) plot from A (see Methods). **C.** Cutoffs (red line) for regions whose UC predominantly correlated with NME or MML in the difference-average plot calculated by looking for local minimums of density (see Methods). **D.** Same cutoffs shown in the original dMML-UC correlation and dNME-UC correlation plot. **E.** Combine MML only and NME only region cutoffs with FDR cutoffs. **F.** Region categorization based on MML cutoff, NME cutoff, and FDR cutoffs. (See Methods)

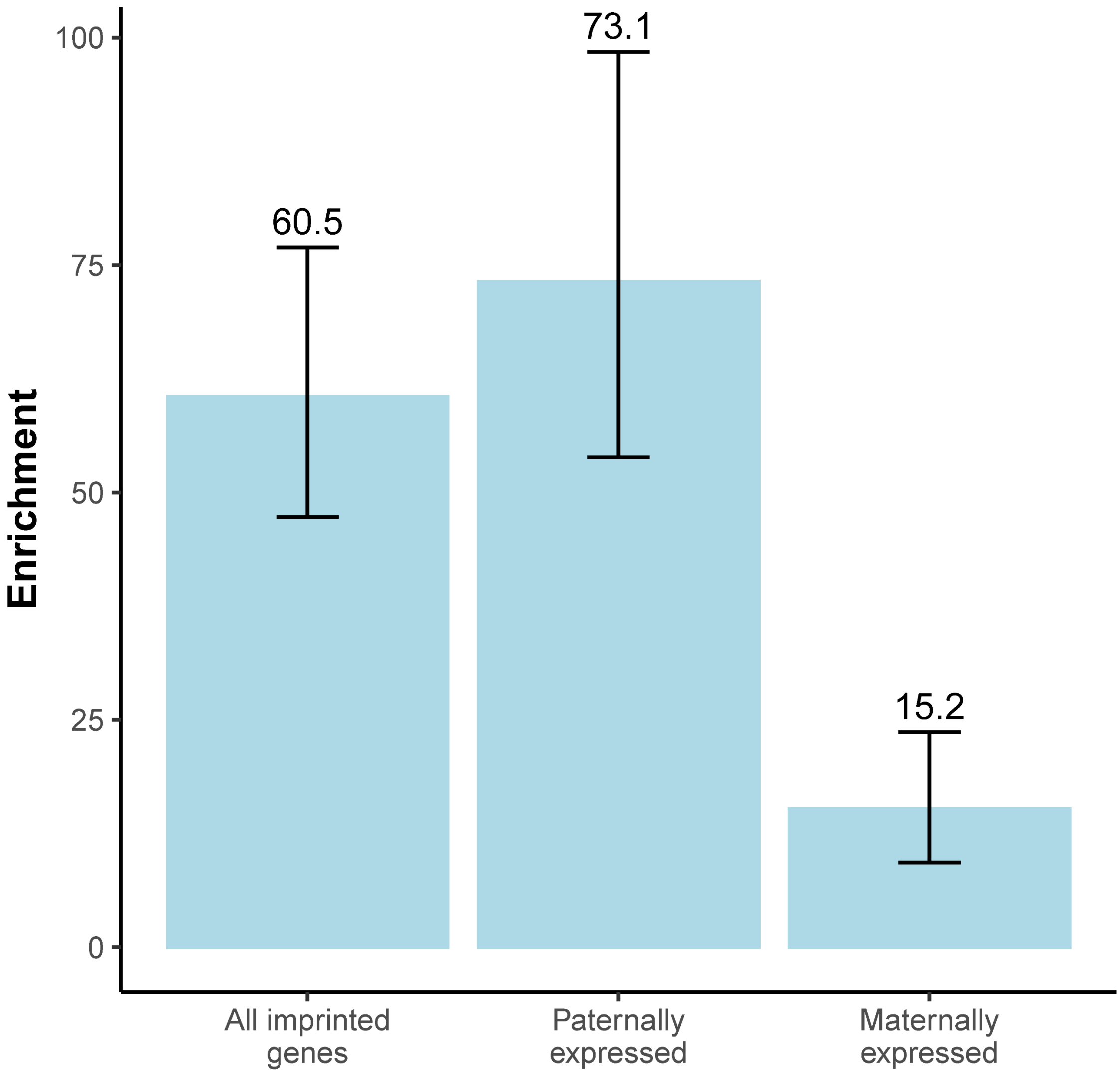


**Fig. S4. The regions with a large mean allelic difference are enriched in the imprinted genes.** The bar plot shows the odds ratio of the regions with a large MML difference in the promoter of imprinted genes. The error bar shows the upper and lower 95% confidence interval of the odds ratio. The regions with a large MML difference are showing high enrichment in the promoters of all imprinted genes.

| Allele1 | Allele2 | Allele 1 has lower NME | Allele 2 has lower NME |
|---|---|---|---|
| GCG | GTG | 1141 | 520 |
| *C* but not GCG | *T* but not GTG | 13362 | 11524 |

Odds Ratio=1.89

P-value<=0.001

Lower CI=1.69

Upper CI=2.11

**Fig S5. Example contingency table to explore trinucleotide context in relation to NME.** Analysis of GCG to GTG in relation to NME. The columns are the number of SNPs showing low NME in each allele. The rows are GCG → GTG trinucleotide and background. The SNPs where the GCG genotype has lower NME than GTG genotype are 1.89 times (95% confidence interval (CI) = [1.69, 2.11]) as high as the odds of having lower NME in the C allele of other C/T SNPs (Fisher's exact test, $P < 2.2 \times 10^{-16}$).

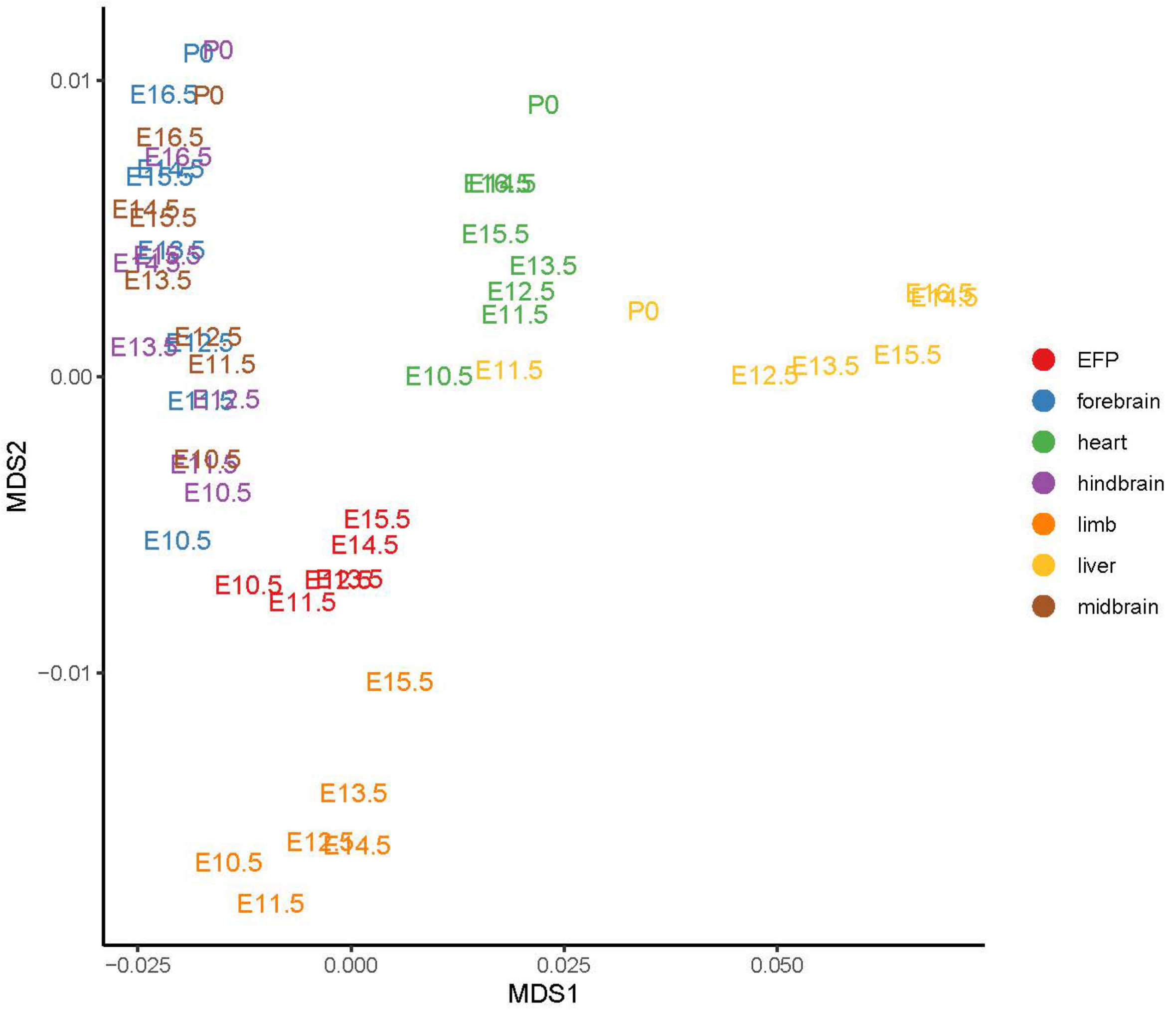


**Fig. S6. MDS plot with postnatal day0 (P0).** Multi-dimensional scaling (MDS) plot of mouse embryonic and postnatal day0 (P0) samples from different tissues and developmental time points. MDS is based on the pairwise distances between samples using uncertainty coefficient (UC). Samples are color-coded by tissues and labeled with development stages. Samples from the same or similar tissues tend to be clustered together. The MDS plot also captures the temporal ordering, with samples from earlier developmental time point closer to a common center (labeled by the red dot) and samples from later time points gradually moving away from the center towards different directions for different tissues. However, the liver P0 sample is between E11.5 and E12.5 and does not follow the developmental trajectory.

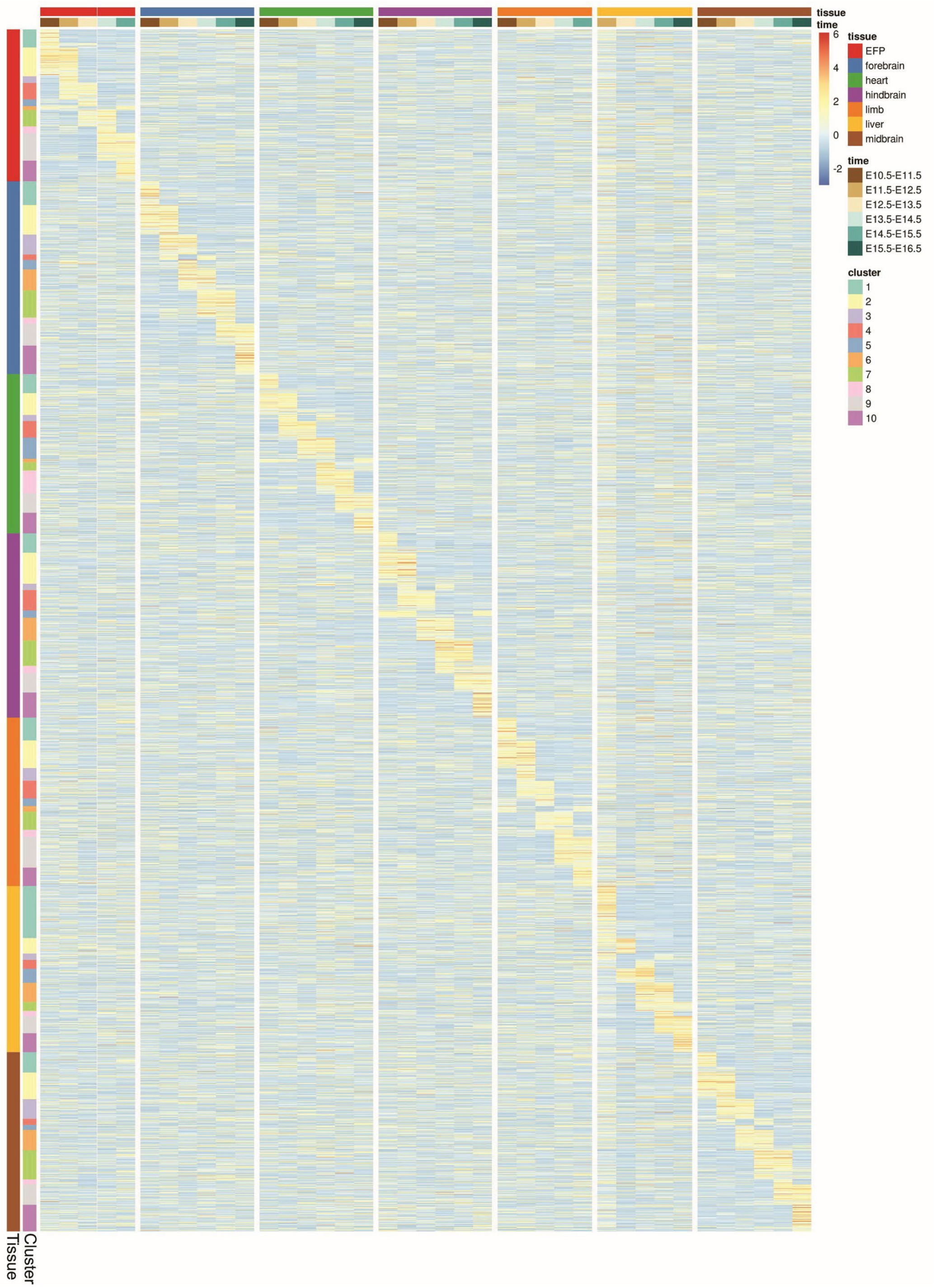


**Fig. S7**. **Clustering analysis of mouse regions with high UC between pairs of time points.** Heatmap of UC values in mouse embryogenesis reveals tissue- and developmental stage-specific DNA methylation landscape changes. Each row is a region with UC > 0.1 in the corresponding tissue, but not limited to that tissue. Each column is a pair of consecutive time points within a tissue. Regions within each tissue are grouped into 10 clusters which were ordered according to which consecutive time points have the largest UC. The clustering result reflected the temporal cascade of methylation landscape changes in that tissue, and also showed that these clusters were tissue-specific. High UC (red) means large methylation change (including mean and entropy changes) and low UC (blue) means small change. The temporal patterns here are tissue-specific, as clustering patterns observed in one tissue are not observed in other tissues.

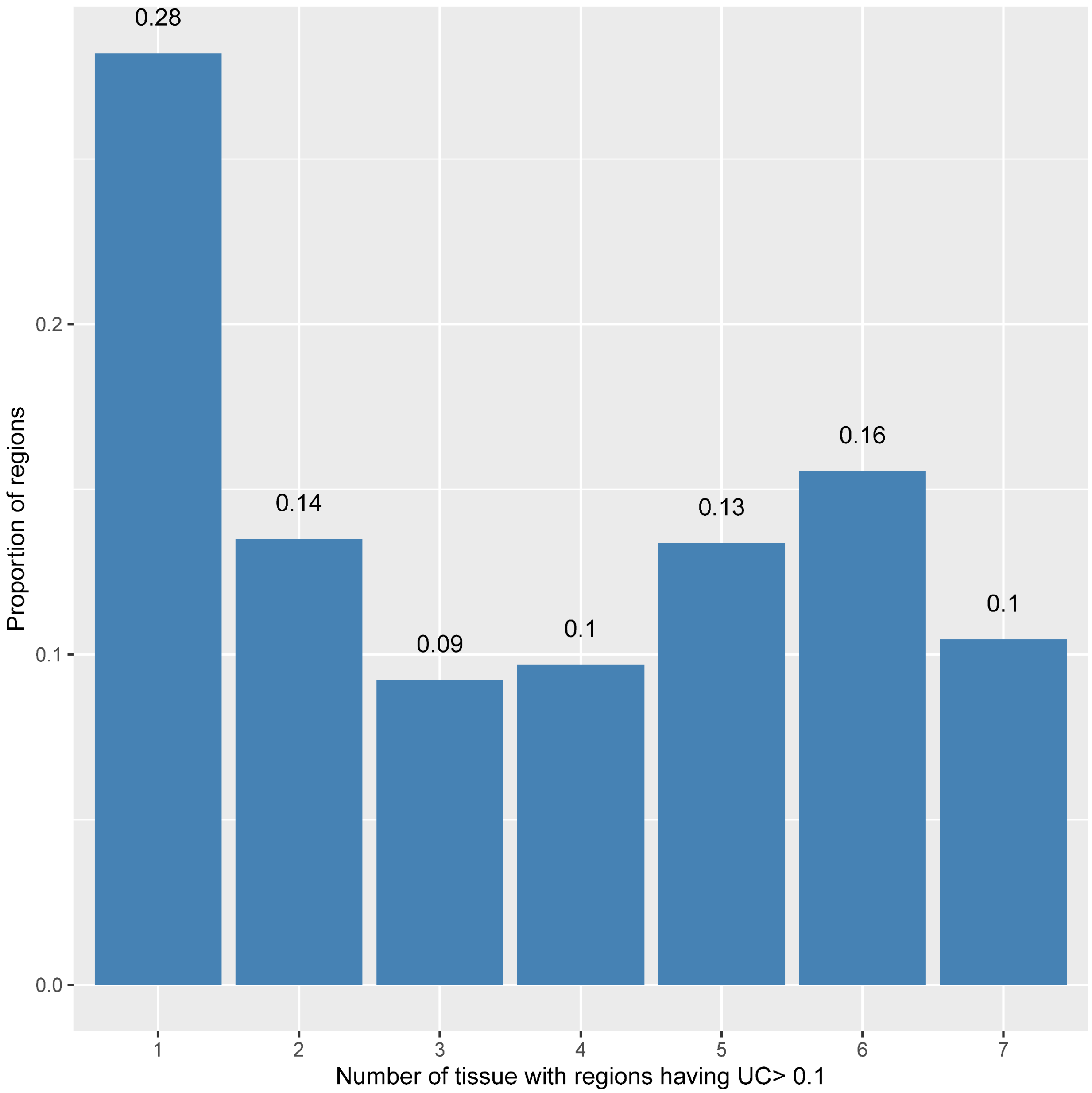


**Fig. S8**. **Proportion of regions where UC > 0.1 between any time point pair, by number of tissues in which it is observed.** The overall distribution was bimodal with the highest peak for unique tissues. Among regions with UC > 0.1 between time points, 28% are present in only one tissue.

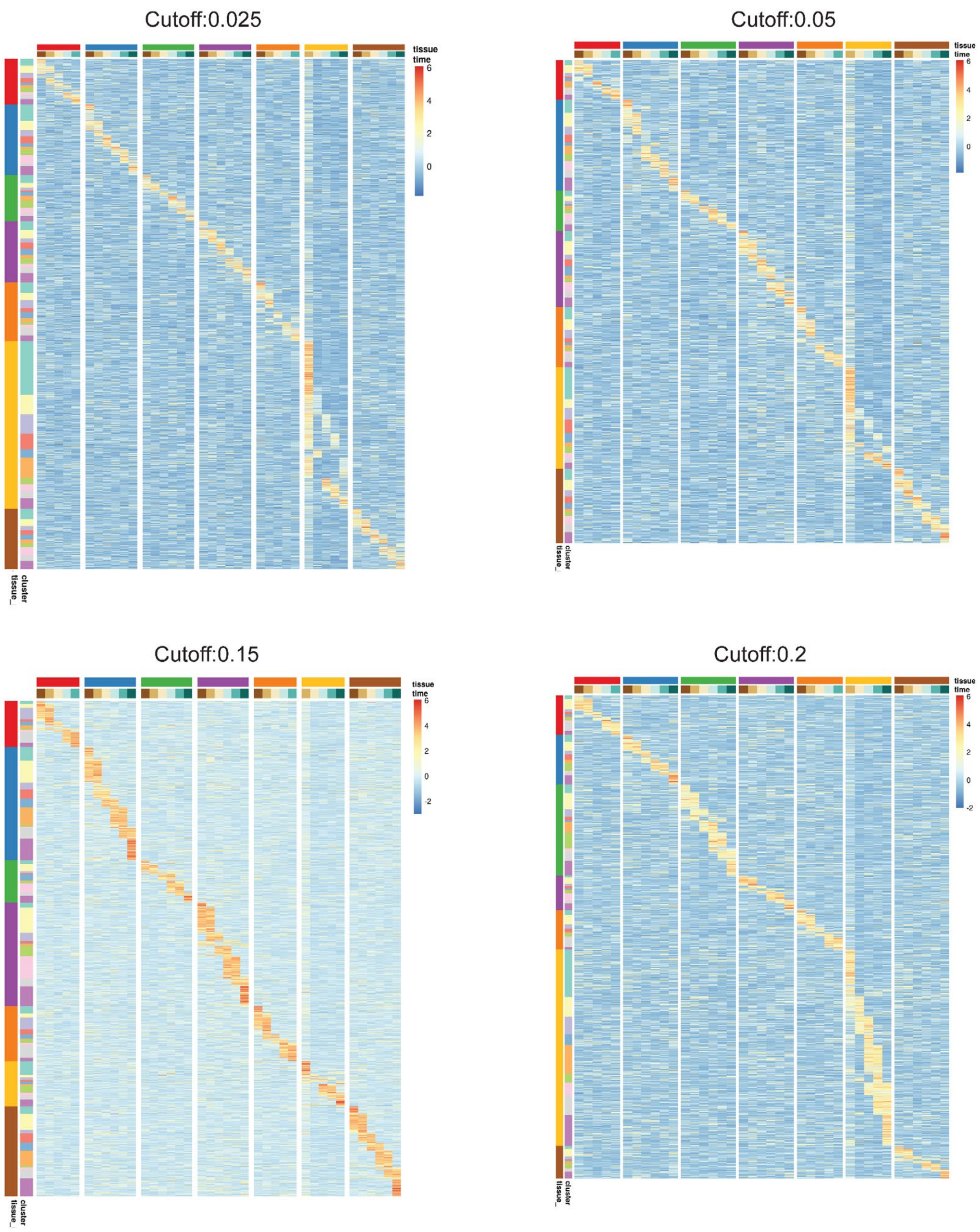


**Fig. S9. Different UC cutoffs generate similar tissue-specific clusters.** Heatmap of UC values in mouse embryogenesis was generated similar to Fig. 4B but using different UC cutoffs. Each column is a pair of consecutive time points within a tissue. Regions within each tissue are grouped into 10 clusters, displayed in the order of their peak UC time, reflecting the temporal cascade of methylation landscape changes in that tissue. High UC (red) means large methylation change (including mean and entropy changes) and low UC (blue) means small change. The temporal patterns here are tissue-specific, as patterns are observed only in one tissue.

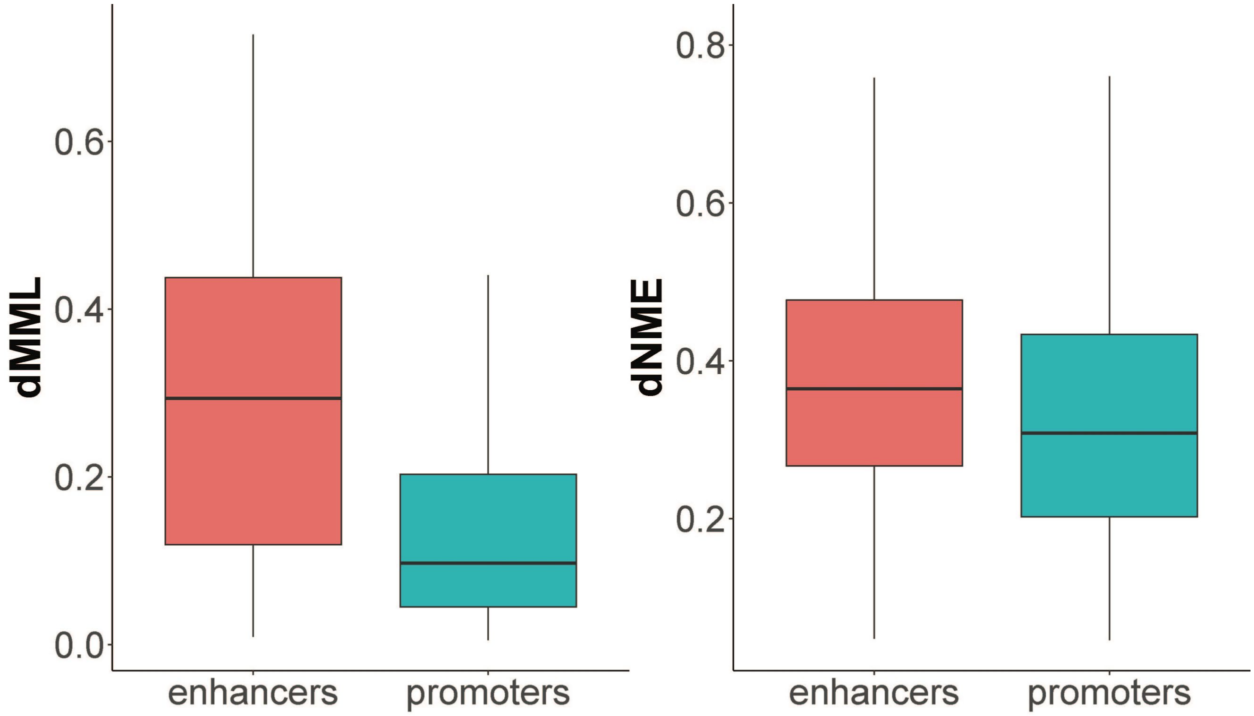


**Fig. S10. Promoters showing lower MML change (dMML) and NME change (dNME) than enhancers.** Boxplots showing dNME levels and dMML levels for enhancer and promoter regions. For each region in each tissue, the dMML and dNME are calculated from all comparisons between adjacent time points, and maximum dMML and dNME for each region are used in the boxplot. The boxplots show that the dNME and dMML are both higher in enhancers than promoters, suggesting more methylation changes at enhancers than at promoters during the development.

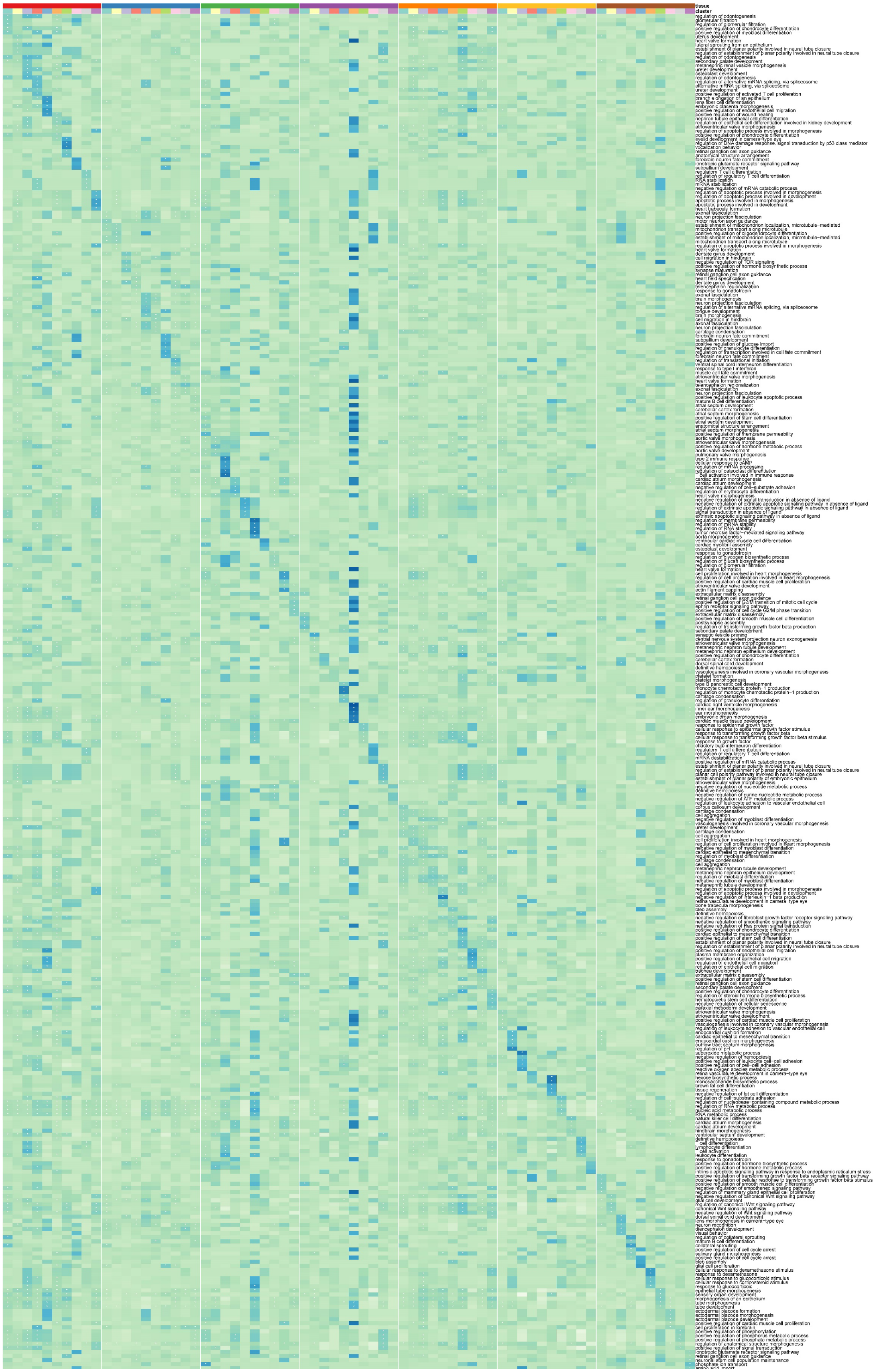

**Fig. S11 Gene ontology (GO) analysis for enhancer regions.** Gene ontology (GO) analysis demonstrates the biological significance of UC-based developmental clustering. For each tissue and region cluster identified in Fig. 4B, enriched biological functions associated with enhancer regions in the cluster are identified and the top 5 GO terms are shown (rows: GO terms; columns: tissues and region clusters; "*": FDR<=0.1). Colors in the heatmap are normalized fold-enrichment of GO terms.

Cutoff:0.025

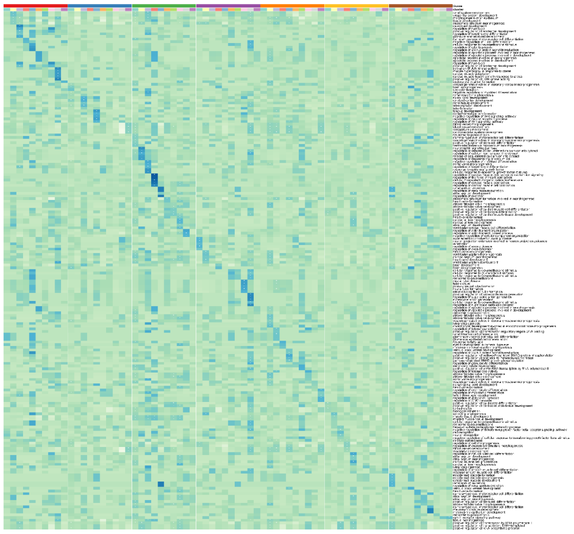

Cutoff:0.05

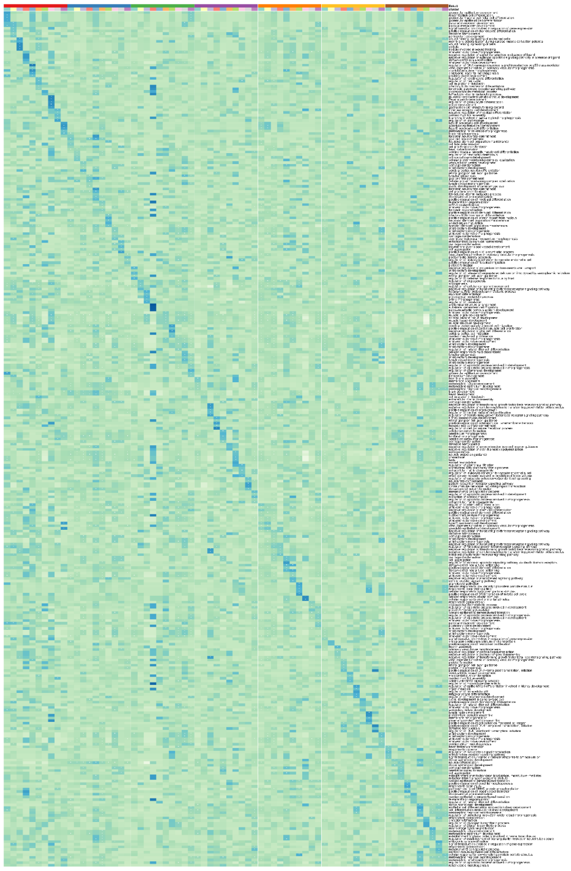

Cutoff:0.15

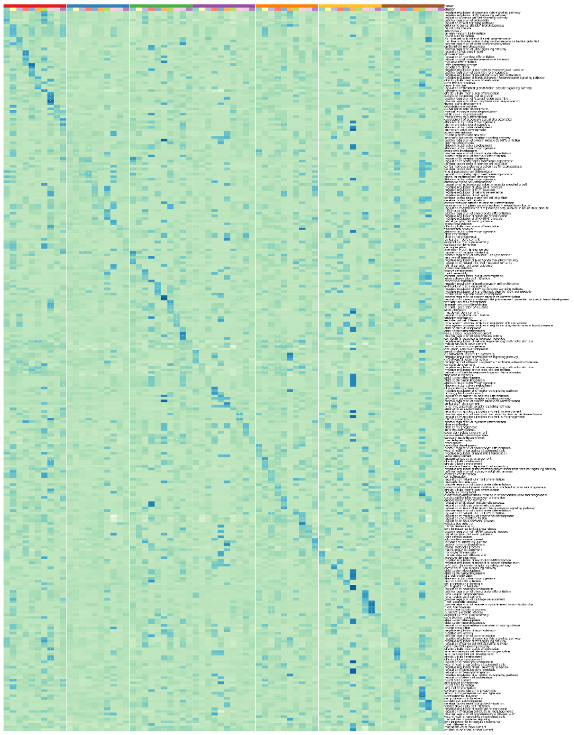

Cutoff:0.2

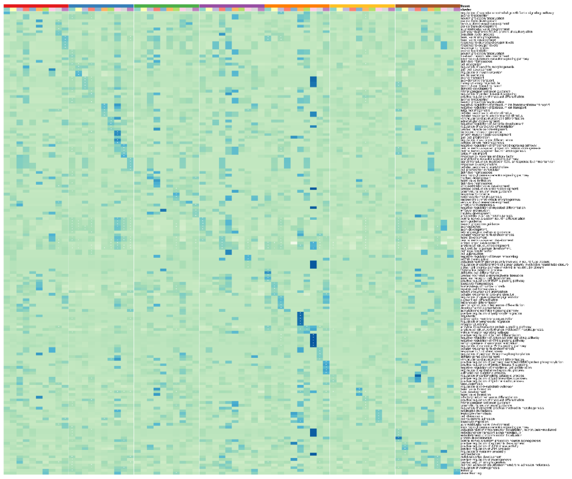

**Fig. S12. Different UC cutoffs generate similar tissue-specific GO results.** Gene ontology (GO) analysis demonstrates the biological significance of UC-based developmental clustering at different UC cutoffs. For each tissue and region cluster identified in Fig. S7, enriched biological functions associated with enhancer regions in the cluster are identified and the top 5 GO terms are shown (rows: GO terms; columns: tissues and region clusters; "*": FDR<=0.1). Colors in the heatmap are normalized fold-enrichment of GO terms. The enriched GO terms reveal tissue-specific functions. Although all cutoffs show tissue-specific GO patterns, the cutoffs of 0.05 and 0.15 show more GO terms and are more tissue-specific. For example, we saw craniofacial suture morphogenesis in EFP, forebrain neuron fate commitment in the forebrain, and artery morphogenesis in the heart using UC cutoff of 0.05 as well as brain morphogenesis in the forebrain, heart valve formation in the heart using UC cutoff of 0.15.

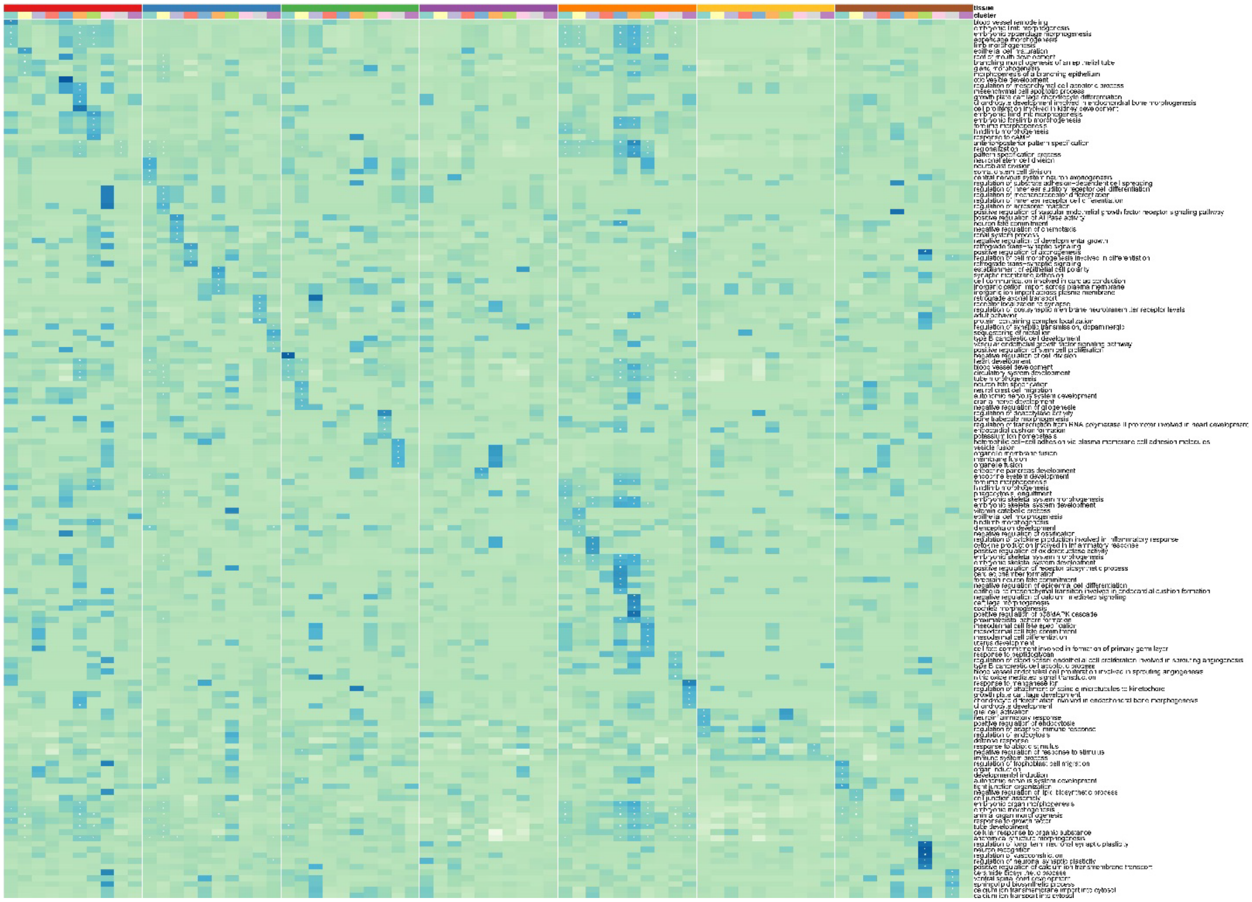

**Fig. S13. Gene ontology (GO) analysis for promoter regions.** Gene ontology (GO) analysis demonstrates the biological significance of UC-based developmental clustering. For each tissue and region cluster identified in Fig. 4B, enriched biological functions associated with promoter regions in the cluster are identified and the top 5 GO terms are shown (rows: GO terms; columns: tissues and region clusters; “*”: FDR ≤ 0.1). Colors in the heatmap are normalized fold-enrichment of GO terms. Many of the terms are related to tissue development such as stem cell differentiation etc. However, very few terms are related to the tissue-specific functions in the development.

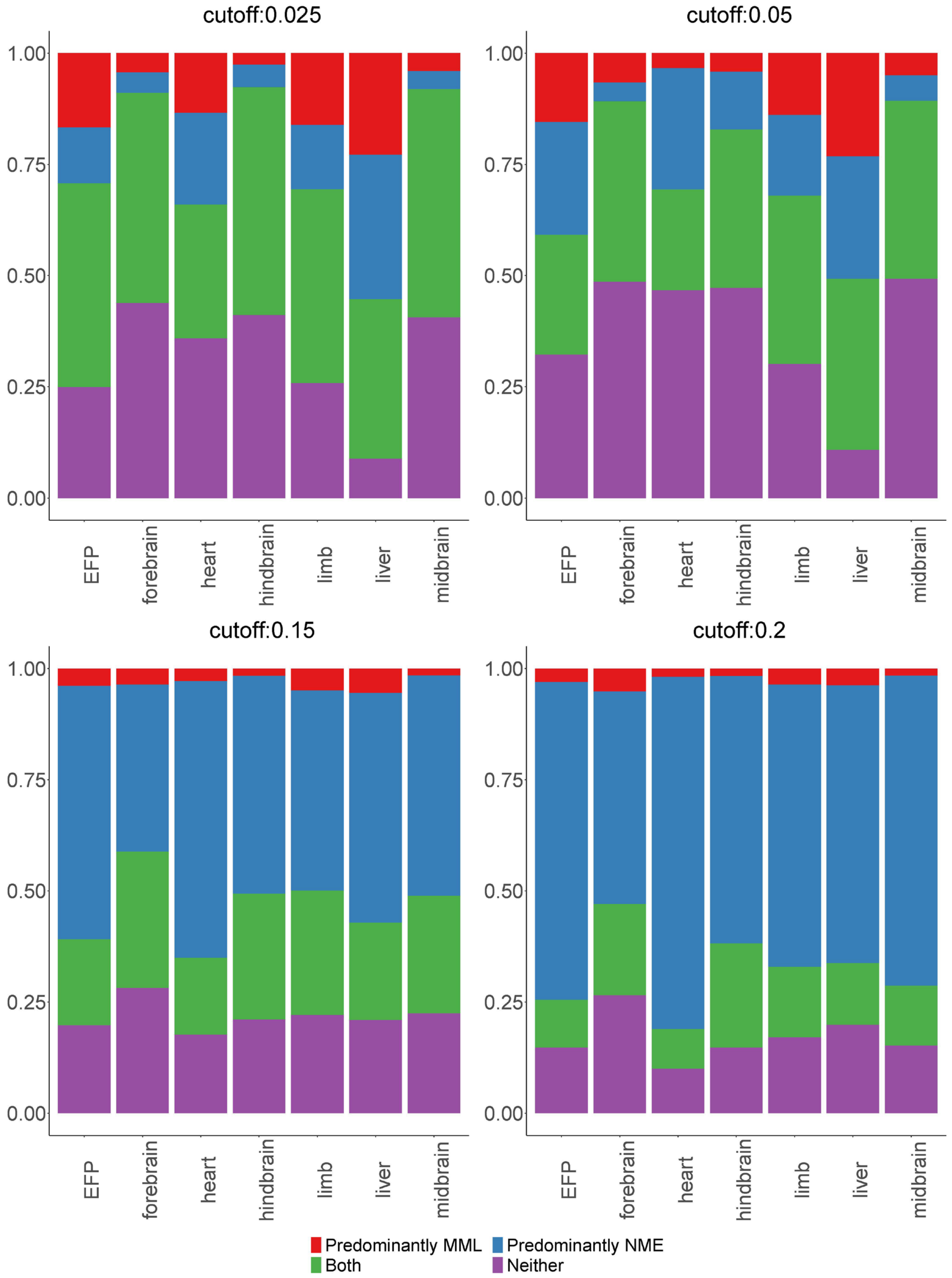


**Fig. S14. Different UC cutoffs have a similar trend in region categorization.** Bar plot showing the relative contribution of MML and NME to the methylation landscape changes as characterized by UC using different UC cutoffs. In each tissue, regions were classified into four categories based on whether a region's UC change across time points was correlated with the change in MML, NME, or both, or uncertain (neither). The proportion of each category among all regions that have high UC in at least one pair of time points is shown. In all different UC cutoffs, there were more regions where UC was predominantly correlated with NME (blue) than the regions where UC was predominantly correlated with MML (MML only, red), suggesting a larger role of entropy than mean in shaping the dynamic changes of methylation landscape.

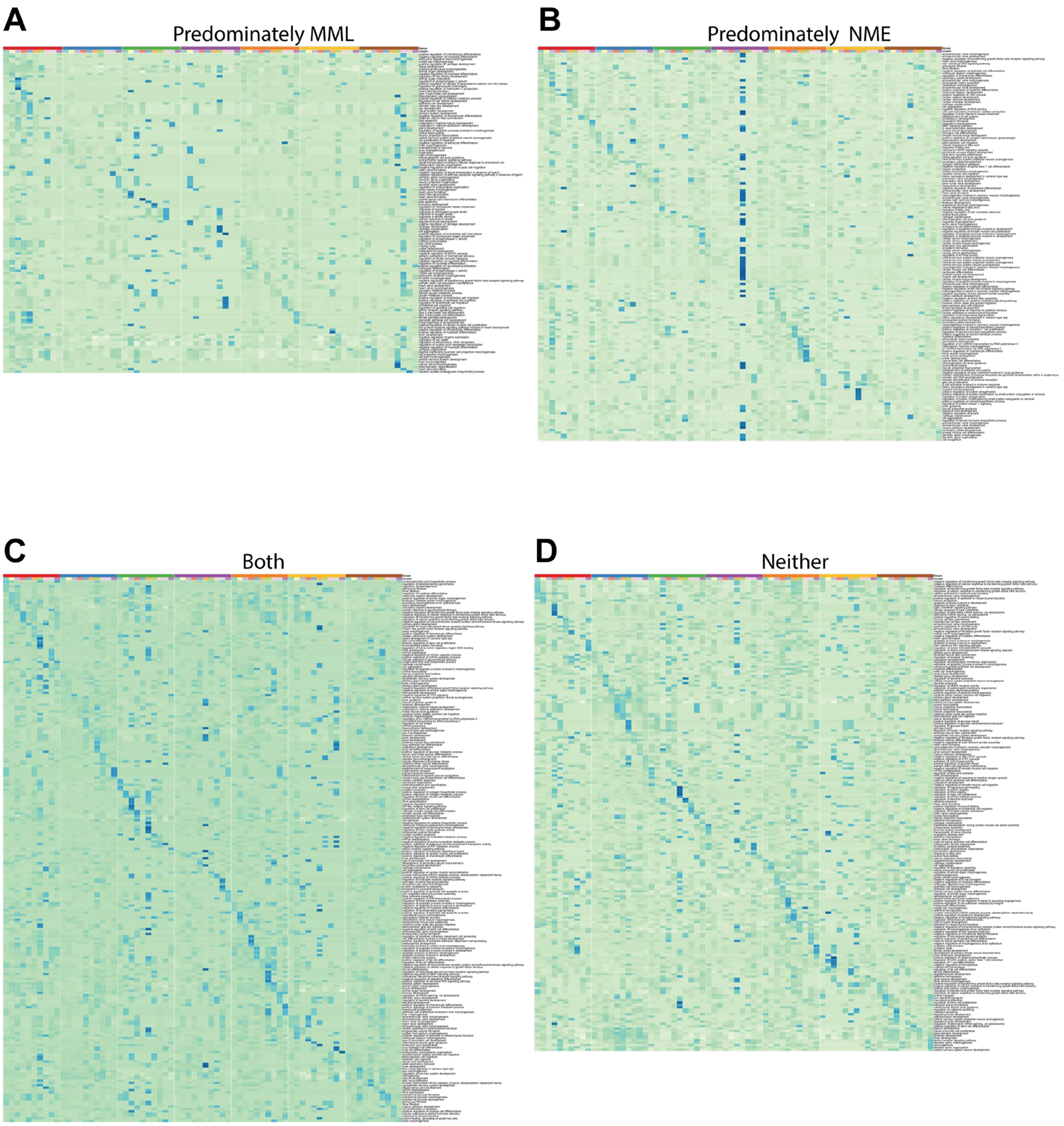


**Fig. S15. Gene ontology (GO) analysis for regions where UC is predominantly correlated with MML, NME, both, or neither.** Gene ontology (GO) analysis demonstrated the biological significance of UC-based developmental clustering. For each tissue and region cluster identified in Fig. 4B, enriched biological functions associated with genes whose UC at enhancers is correlated with MML, NME, both or neither in the cluster were identified and the top 5 GO terms are shown (rows: GO terms; columns: tissues and region clusters; "*": FDR<=0.1). Colors in the heatmap were normalized fold-enrichment of GO terms. **A.** The UC in those regions was predominantly correlated with MML. Many of the terms were related to tissue development such as stem cell differentiation etc. However, very few terms were related to the tissue-specific functions in the development. Some tissues, such as the hindbrain, did not show significant terms. **B.** The UC in those regions was predominantly correlated with NME. This was showing more terms than MML only regions suggesting NME also contain information about tissue development. We also observe tissue-specific terms such as atrioventricular valve development in the heart. **C.** The UC in those regions was predominantly correlated with both. **D.** The UC in those regions was correlated with neither NME nor MML.

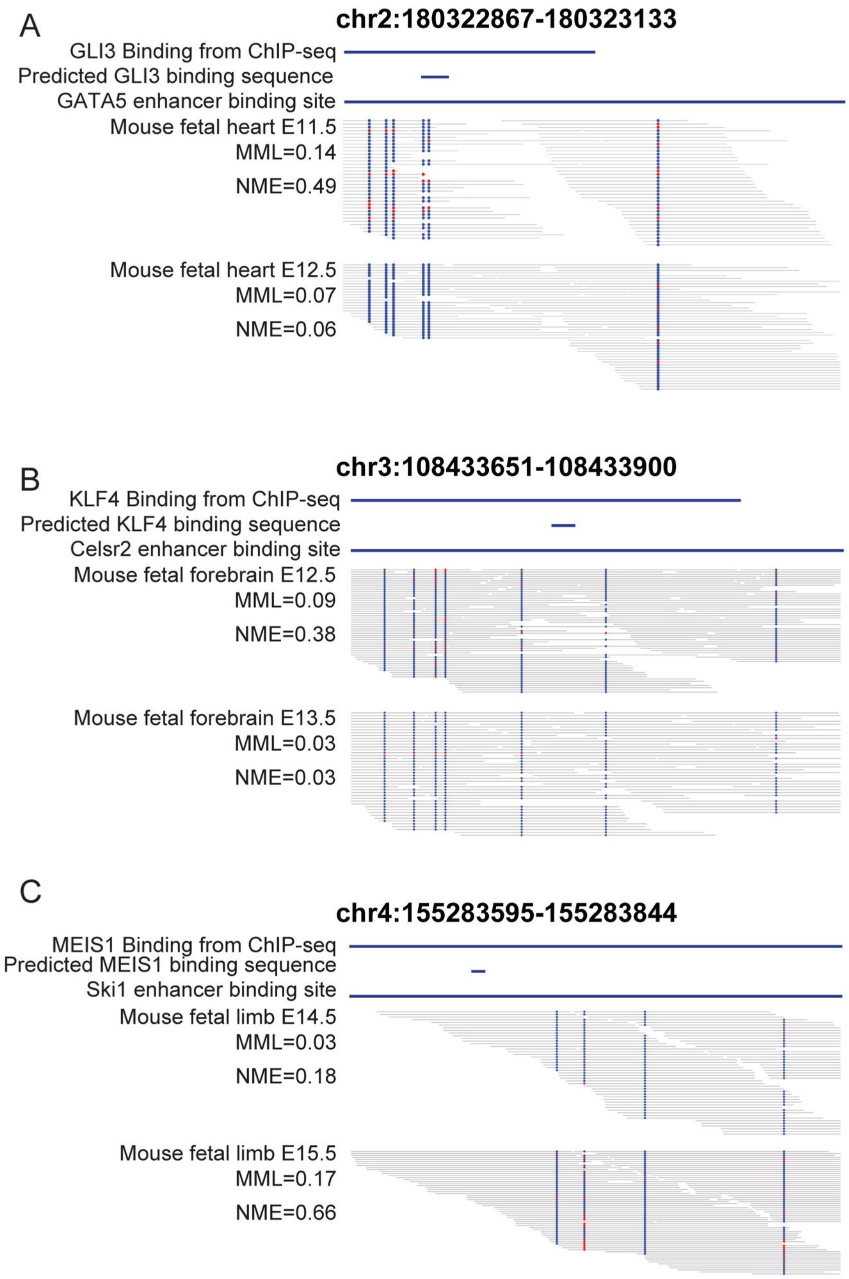


**Fig. S16. Examples showing motif binding sites having large NME change during development.** Example regions that are enhancers of selected genes and the binding sites of the transcription factors that are enriched in NME only regions for the corresponding tissue in Fig 6A, B, C. Each continuous grey line is a read. Each circle is a CpG site. The blue circle is unmethylated CpG and the red circle is methylated CpG. The potential energy landscape is modeled on each read, i.e., each grey line. If two or more grey lines are in the same row, the potential energy landscape is modeled separately for each grey line. Zoom in to view the methylation state at each read. **A.** The region in mouse fetal heart E11.5 and E12.5 is an enhancer of the *Gata5* gene and a GLI3 binding site from the ChIP-seq experiment [1]. The first 2 rows are the motif binding site from the ChIP-seq experiment and the predicted motif binding site for GLI3 from the sequence. The third row is the enhancer-binding site. The first block of lines and circles are the reads from WGBS data of mouse fetal heart at E11.5. The second block is the WGBS reads from mouse fetal heart E12.5. We found the methylation pattern in the mouse fetal heart E12.5 is much less random than the methylation pattern in mouse fetal heart E11.5. **B.** The region in the mouse fetal forebrain. The organization is the same as A. We saw the methylation pattern is highly organized in mouse fetal forebrain E13.5 suggesting low NME at E13.5. Consistent with the higher NME in E12.5, the methylation pattern is more variable in E12.5 than E13.5. **C.** The region in the mouse fetal limb. The organization is the same as A. We saw the methylation pattern is less organized in mouse fetal limb E15.5 than E14.5.

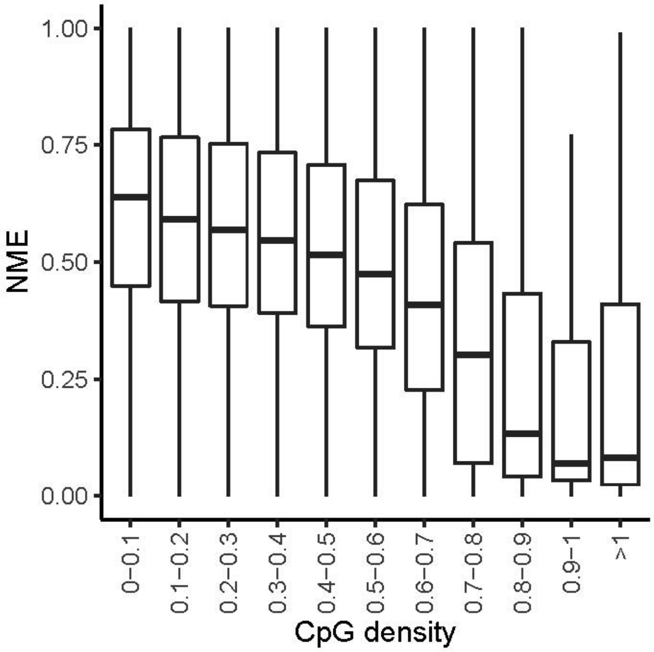


**Fig. S17 Decrease in NME as CpG density increase in the mouse.** NME regardless of SNPs, of genomic regions, stratified based on CpG density. The CpG density is defined as the ratio of the observed CpG number to the expected CpG number in each region. As CpG density increases, the NME decreases.

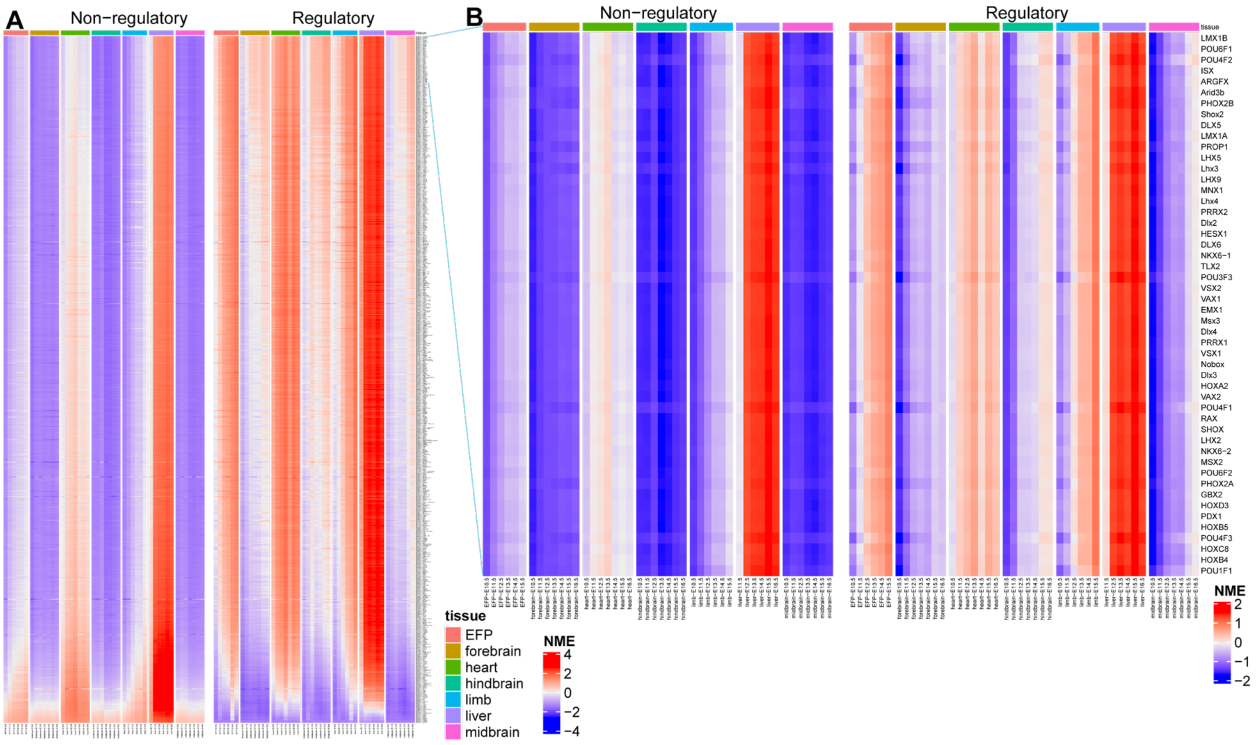


**Fig. S18. Transcription factor (TF) binding motif sites have higher normalized methylation entropy (NME) at regulatory DNA than non-regulatory DNA in the mouse. A.** Heatmaps comparing NME between regulatory DNA (right panel) and non-regulatory DNA (left panel) for the 736 motifs in mouse from JASPAR database. Each row corresponds to a TF binding motif and each column corresponds to a WGBS sample. For each motif and in each WGBS sample, the median NME across all regulatory (right panel) or non-regulatory (left panel) DNA with the motif is shown. For visualization, NME values within each row were standardized across regulatory and non-regulatory DNA to have zero mean and unit SD (red: high NME; blue: low NME). **B.** Heatmaps comparing NME between regulatory DNA (right panel) and non-regulatory DNA (left panel) for the top 50 motifs in **A** with the largest difference between regulatory and non-regulatory DNA.

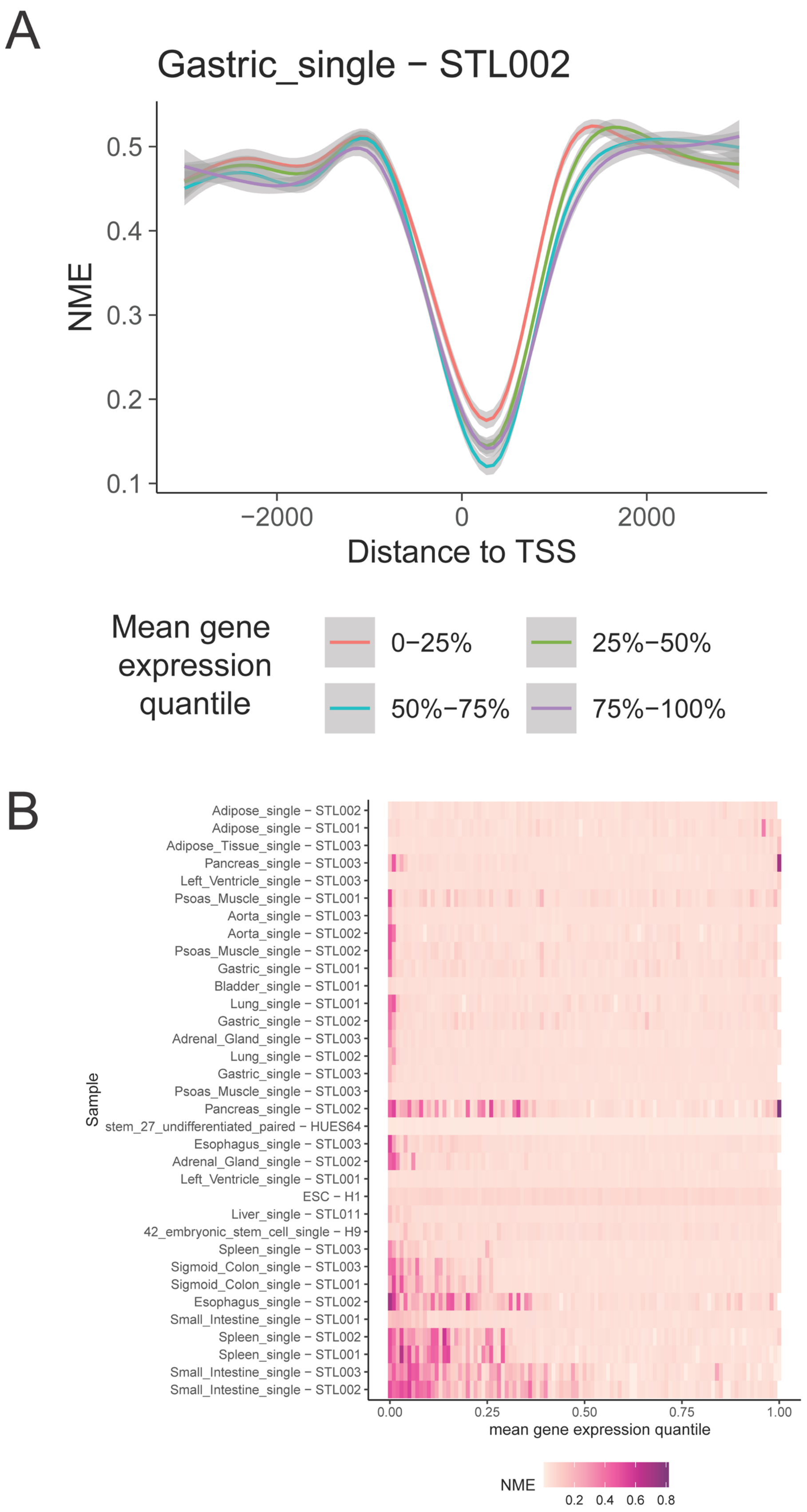
A
Gastric_single − STL002
NME
0.5
0.4
0.3
0.2
0.1
−2000
0
2000
Distance to TSS
Mean gene expression quantile
0−25%
25%−50%
50%−75%
75%−100%
B
Sample
Adipose_single − STL002
Adipose_single − STL001
Adipose_Tissue_single − STL003
Pancreas_single − STL003
Left_Ventricle_single − STL003
Psoas_Muscle_single − STL001
Aorta_single − STL003
Aorta_single − STL002
Psoas_Muscle_single − STL002
Gastric_single − STL001
Bladder_single − STL001
Lung_single − STL001
Gastric_single − STL002
Adrenal_Gland_single − STL003
Lung_single − STL002
Gastric_single − STL003
Psoas_Muscle_single − STL003
Pancreas_single − STL002
stem_27_undifferentiated_paired − HUES64
Esophagus_single − STL003
Adrenal_Gland_single − STL002
Left_Ventricle_single − STL001
ESC − H1
Liver_single − STL011
42_embryonic_stem_cell_single − H9
Spleen_single − STL003
Sigmoid_Colon_single − STL003
Sigmoid_Colon_single − STL001
Esophagus_single − STL002
Small_Intestine_single − STL001
Spleen_single − STL002
Spleen_single − STL001
Small_Intestine_single − STL003
Small_Intestine_single − STL002
0.00
0.25
0.50
0.75
1.00
mean gene expression quantile
NME
0.2
0.4
0.6
0.8

**Fig. S19. Mean gene expression is not positively associated with NME. A.** Association between NME surrounding transcription start sites (TSS) and mean gene expression. The example shows the averaged NME near TSS at different mean gene expression quantiles. **B.** Heatmap showing NME values within +500 bp of TSS in the sample. In each sample, genes are stratified based on their mean gene expression using quantiles of mean gene expression (X-axis), and the heatmap shows the average NME of all genes in each stratum. For most tissues, we did not observe increased NME (from pink to dark red) as the mean gene expression quantile increases, indicating that mean gene expression is not positively correlated with NME in regions surrounding TSS.

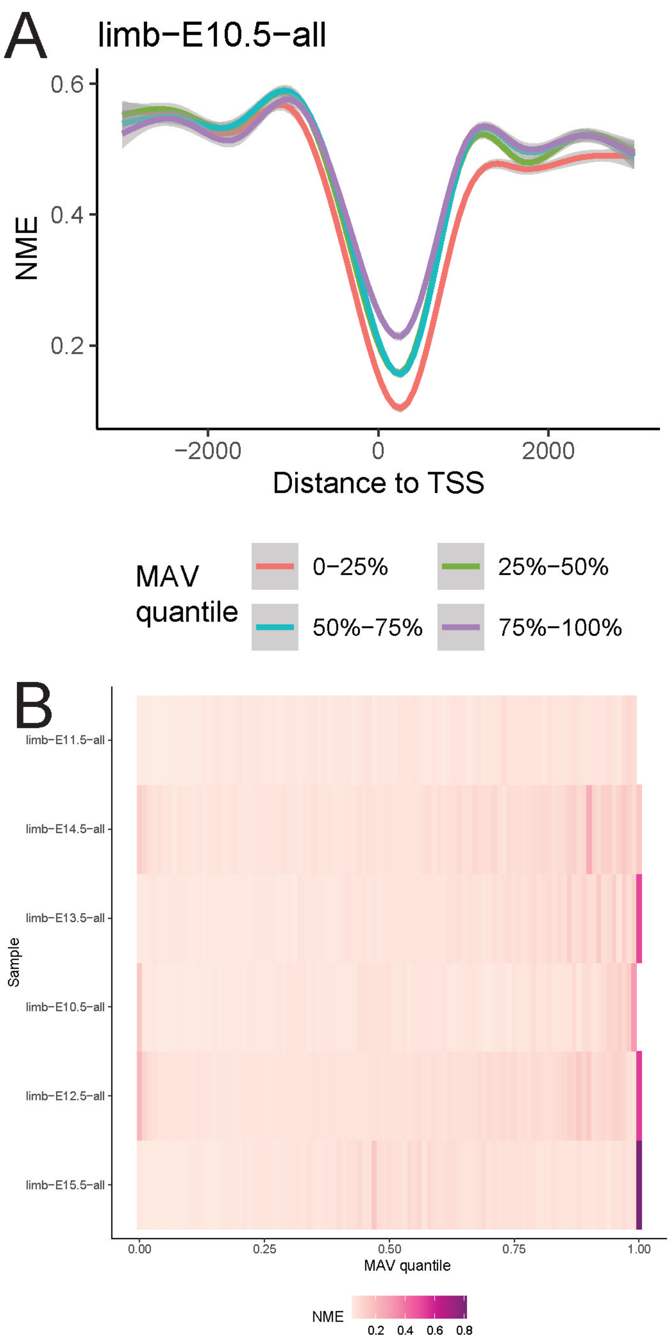


**Fig. S20. NME is positively correlated with MAV in the mouse. A.** Association between NME surrounding transcription start sites (TSS) and gene expression variability. Genes were stratified based on quartiles of their mean-adjusted expression variability (MAV). The example shows the averaged NME near TSS is lower for the gene with lower MAV. For each stratum, the average NME surrounding TSS across all genes in the stratum is shown. The genes in the lowest MAV quantile (red line) have the lowest NME near their TSS. The genes in the highest MAV quantile (purple line) have much higher NME near their TSS. **B.** Heatmap showing NME values within +500 bp of TSS in different stages during mouse fetal limb development. In each sample, genes are stratified based on their expression variability using quantiles of MAV (X-axis), and the heatmap shows the average NME of all genes in each stratum. For most tissues, we observe increased NME (from pink to dark red) as the MAV quantile increases, indicating that gene expression variability (MAV) is correlated with NME in regions surrounding genes' TSS.

**Dataset S1. Transcription factors whose motifs are associated with MML in human.** This table is a list of transcription factors whose motifs are associated with MML. The column labeled "proportion" is the proportion of motif sites for that motif that binds to a higher MML allele or lower MML allele. There are significantly more motifs that are associated with low MML than motifs that are associated with high MML

**Dataset S2. Transcription factors whose motifs are associated with NME in human.** This table is a list of transcription factors whose motifs are associated with NME. The column labeled "proportion" is the proportion of motif sites for that motif that bind to a higher NME allele or lower NME allele. There are significantly more motifs that are associated with high NME motifs that are associated with low NME

**Dataset S3. Transcription factors whose motifs have high NME but not low MML.** This table is a list of transcription factors whose motifs with high NME but not low MML. We found 55 transcription factors that are only associated with high NME. The column labeled "proportion" is the proportion of motif sites for that motif that bind to a higher NME allele or lower NME allele.

**Dataset S4 NME difference of regulatory DNA minus non-regulatory DNA at motif binding sites in human.** This shows the difference of NME regardless of allele between motif sites at regulatory DNA and motif sites at non-regulatory DNA for 630 motifs based on all genomic regions in human. P-values are obtained from Wilcoxon signed-rank tests for comparing the NME between regulatory and non-regulatory DNA for each motif. P-values are converted to FDRs using the Benjamini–Hochberg procedure.

**Dataset S5 Enriched pathway in transcription factor whose motifs show the largest difference in NME between regulatory and non-regulatory DNA** This table shows the GO terms from GOrilla analysis using all transcription factors ranked by their difference. The gene columns show the genes in each term that overlap with the 50 genes showing the highest NME difference as in fig S2B.

**Dataset S6 Transcription factor (TF) binding motif sites have higher normalized methylation entropy (NME) at regulatory DNA than non-regulatory DNA.** This table shows the difference of NME, regardless of allele, between motif sites at regulatory DNA and motif sites at non-regulatory DNA for 630 motifs based on all genomic regions in human. P-values are obtained from Wilcoxon signed-rank tests for comparing the NME between regulatory and non-regulatory DNA for each motif. P-values are converted to FDRs using the Benjamini–Hochberg procedure.

**Dataset S7. List of GO terms for genes whose enhancers have high UC changes.** This is a list of GO terms for each tissue using all enhancer regions. Each tab is a tissue. We observe the GO terms ($2^{nd}$ column) are enriched in tissue-specific GO terms.

**Dataset S8. List of GO terms for genes whose UC at enhancers was correlated predominantly with NME.** This is a list of GO terms for each tissue using all enhancer regions whose UC is predominantly correlated with NME. Each tab is a tissue. We observed GO terms specific to the tissue, such as “atrioventricular valve development” in the heart.

**Dataset S9. List of GO terms for genes whose UC at enhancers was correlated predominantly with MML.** This is a list of GO terms for each tissue using all enhancer regions whose UC was predominantly correlated with MML. Each tab is a tissue. We observed fewer terms enriched in the heart compared with Dataset S8 and no terms in the hindbrain.

**Dataset S10. List of GO terms for genes whose UC at enhancers was correlated with both NME and MML.** This is a list of GO terms for each tissue using all enhancers whose UC was correlated with both NME and MML. Each tab is a tissue. We observed many tissue-specific terms, but they are different than the ones we observed in Dataset S8. For example, the top term in the heart is “muscle cell development”.

**Dataset S11. List of GO terms for genes whose UC at enhancers was independent of both NME and MML.** This is a list of GO terms for each tissue using all enhancer regions whose UC is independent of both MML and NME. Each tab is a tissue. There are some GO terms enriched in the tissue, such as “regulation of nitrogen compound metabolic process” and other metabolism-related terms in the heart suggesting there are methylation landscape changes that cannot be explained by either MML or NME.

**Dataset S12. Motifs enriched in regions correlated predominantly with MML or NME in the mouse.** This table shows the lists of motifs that are enriched in NME only regions and MML only regions in different mouse tissues. Residual represents the residual of a motif to the fitted linear regression line. Log_OR represents the log odds ratio for the enrichment of a motif in each type of region (i.e., correlated predominantly with NME or MML). Normalized_log_OR represents the normalized log odds ratio. FDR represents the false discovery rate for the motif enrichment test.

**Dataset S13. List of genes whose UC at enhancers was correlated predominantly with NME** This is a list of genes in the enriched GO terms whose UC at enhancer regions was correlated predominantly with NME and overlaps motif binding sites for the motifs associated with methylation entropy. The first column is the tissue in which we found the genes/motifs. The second column is the predicted transcription factor motif that overlaps with the enhancer. The third column is the continuous stages where we find the maximum UC. The fourth column is the genes regulated by enhancers. The fifth and seventh columns are the dNME and dMML at the stages where we find the maximum UC. The sixth and eighth columns are the correlation between dNME or dMML and UC. The column labeled “PMID” shows the reference PMID that is associated with that gene and tissue. We find many genes that are important in corresponding tissue.

**Dataset S14. List of genes whose enhancers are in regions where UC was predominatly corerlated with MML and containing motifs.** This is a list of genes in the enriched GO terms whose enhancer regions regions where UC was predominatly corerlated with MML and overlaps motif binding sites for the motifs associated with methylation entropy. The first column is the tissue in which we found the genes/motifs. The second column is the predicted transcription factor motif that overlaps with the enhancer. The third column is the continuous stages where we find the maximum UC. The fourth column is the genes regulated by enhancers. The fifth and seventh columns are the dNME and dMML at the stages where we find the maximum UC. The sixth and eighth columns are the correlation between dNME or dMML and UC. The column labeled "PMID" shows the reference PMID that is associated with that gene and tissue. We find many genes that are important in corresponding tissue.

**Dataset S15. NME difference of regulatory DNA minus non-regulatory DNA at motif binding sites in the mouse.** This shows the difference of NME, regardless of allele, between motif sites at regulatory DNA and motif sites at non-regulatory DNA for 736 transcription factor motifs from genomic regions in the mouse. P-values are obtained from Wilcoxon signed-rank tests for comparing the NME between regulatory and non-regulatory DNA for each motif. P-values are converted to FDRs using the Benjamini–Hochberg procedure.

**Dataset S16. MAE enrichment in regions with strong MML or NME difference.** This shows the contingency table to calculate the enrichment of the regions with strong MML difference or NME difference in the promoters for MAE genes. There is no enrichment of the regions with strong NME difference in the promoter of MAE genes but enrichment of the regions with strong MML difference.

**Dataset S16. MAE enrichment in regions with strong MML or NME difference.** This shows the contingency table to calculate the enrichment of the regions with strong MML difference or NME difference in the promoters for MAE genes. There is no enrichment of the regions with strong NME difference in the promoter of MAE genes but enrichment of the regions with strong MML difference
